# An ER CREC family protein regulates the egress proteolytic cascade in malaria parasites

**DOI:** 10.1101/457481

**Authors:** Manuel A. Fierro, Beejan Asady, Carrie F. Brooks, David W. Cobb, Alejandra Villegas, Silvia N.J. Moreno, Vasant Muralidharan

**Author notes:** Address correspondence to Vasant Muralidharan.

## Abstract

The endoplasmic reticulum (ER) is thought to play an essential role during egress of malaria parasites because the ER is assumed to be the calcium (Ca^2+^) signaling hub and required for biogenesis of egress-related organelles. However, no proteins localized to the parasite ER have been shown to play a role in egress of malaria parasites. In this study, we generated conditional mutants of the *Plasmodium falciparum* <u>E</u>ndoplasmic <u>R</u>eticulum-resident <u>C</u>alcium-binding protein (PfERC), a member of the CREC family. Knockdown of PfERC shows that this gene is essential for asexual growth of *P. falciparum*. Analysis of the intraerythrocytic lifecycle revealed that PfERC is essential for parasite egress but not required for protein trafficking or Ca^2+^ storage. We found that PfERC knockdown prevents the rupture of the parasitophorous vacuole membrane. This is because PfERC knockdown inhibited the proteolytic maturation of the subtilisin-like serine protease, SUB1. Using double mutant parasites, we show that PfERC is required for the proteolytic maturation of the essential aspartic protease, Plasmepsin X, which cleaves SUB1. Further, we show that processing of substrates downstream of the proteolytic cascade is inhibited by PfERC knockdown. Thus, these data establish the ER-resident CREC family protein, PfERC, as a key early regulator of the egress proteolytic cascade of malaria parasites.

## Introduction

Members of the phylum *Apicomplexa* are responsible for severe human diseases such as malaria, toxoplasmosis, and cryptosporidiosis. Together, this group of obligate intracellular parasites causes several hundred million infections every year and remains one of the major drivers of infant mortality (1–4). In fact, malaria results in nearly half a million deaths each year and most of the mortality is attributed to one species, *Plasmodium falciparum*. All the clinical symptoms of malaria are directly correlated to the asexual lifecycle of malaria parasites within the host red blood cells.

The egress and subsequent invasion of daughter parasites into the host cells are essential for the propagation of apicomplexan parasites. Both egress and invasion are ordered and essential processes which are regulated by signaling pathways dependent upon the second messengers, cGMP and Ca^2+^ (5–9). Upon invading the host cell, the parasites create and reside within a host-derived vacuole called the parasitophorous vacuole (PV). Within this vacuole, the parasites grow and divide into daughter cells, which must egress from the host cell to complete the life cycle. This event depends upon the generation of specific organelles that are populated by proteases as well as invasion ligands. These organelles are released via timed exocytosis, which is regulated by signaling pathways dependent upon the second messengers, cGMP and Ca^2+^ (5–9). For example, inhibition of the cGMP dependent protein kinase (PKG) activity blocks egress (6, 10, 11). Ca^2+^-signaling also induces egress although it is uncertain whether this pathway works downstream (7) or synergistically with cGMP signaling (6, 12, 13). The end result of these signaling pathways is thought to be the exocytosis of egress-related vesicles such as exonemes. Exonemes are populated by proteases such as the aspartic protease, Plasmepsin X (PMX) and the serine protease, Subtilisin 1 (SUB1) and the release of mature PMX and SUB1 into the PV commits the parasites for egress resulting in the rapid (∼10 minutes) breakdown of the parasitophorous vacuole membrane (PVM) and the RBC membrane (RBCM)(14, 15).

In malaria parasites, the proteases present in egress-related vesicles require proteolytic cleavage to be activated (14, 16, 17). For example, SUB1 undergoes two cleavage events. First, the zymogen undergoes Ca^2+^ dependent autoprocessing in the ER (18, 19) and then, it is cleaved again by PMX (16, 17). In turn, PMX itself is processed from a 67kDa zymogen to a 45kDa active protease (16, 17). Unlike most aspartic proteases, PMX does not undergo auto proteolysis for maturation as PMX inhibitors are unable to prevent its cleavage into the 45kDa form (16, 17). This suggests that there are as yet unknown factors that regulate the maturation of these key egress proteases. However, there are no obvious candidates in the genome that could regulate these activities.

In this work, we focused on the parasite endoplasmic reticulum (ER) because proteins localized to this organelle are thought to play a key role in egress of daughter merozoites. Their putative functions during this lifecycle stage include the biogenesis of specific egress-related organelles, transporting proteins to these organelles, and propagating Ca^2+^ signals essential for egress (20, 21). However, none of the proteins responsible for these functions during egress of apicomplexan parasites have been identified. One potential candidate is the ER-resident calcium binding protein PfERC (PF3D7_1108600). In malaria parasites, PfERC is the only protein with identifiable Ca^2+^-binding domains localized to the ER and it is capable of binding Ca^2+^ (22). However, the biological function of PfERC is unknown. To address this, we used CRISPR/Cas9 based gene editing approach to generate conditional mutants of PfERC as well as double conditional mutants of PfERC and PMX. These mutants allowed us to determine that this ER-resident protein controls the nested proteolytic cascade in *P. falciparum* that regulates the egress of malaria parasites from human RBCs.

## Results

### PfERC is an CREC family protein localized in the ER

PfERC is a protein related to the CREC (<u>C</u>alumenin, <u>R</u>eticulocalbin 1 and 3, <u>E</u>RC-55, <u>C</u>ab-45) family of proteins, which are characterized by the presence of multiple EF-hands and localization in various parts of the secretory pathway (23, 24) (Figure 1A and Supplementary Figure 1). PfERC contains a signal peptide, multiple EF-hands, and an ER-retention signal (Figure 1A). The domain structure of PfERC is homologous to other members of the CREC family of proteins (Figure 1A and Supplementary Figure 1). However, PfERC differs from its mammalian homologs in that it only contains 5 predicted EF-hands although a 6^th^ degenerate EF-hand (residues 314-325) may be present in its extended C-terminus (Supplementary Figure 1) (22). Various roles have been attributed to CREC members including Ca^2+^ signaling and homeostasis, and one member, RCN3, has been shown to interact with the subtilisin-like peptidase, PACE4, though the functional significance of this interaction is unknown (23, 25). As PfERC expression peaks during early schizont stage parasites, we hypothesized that PfERC is required for egress of daughter parasites during this terminal stage of the asexual lifecycle (22, 26) (Figure 1A).

**Figure 1:**
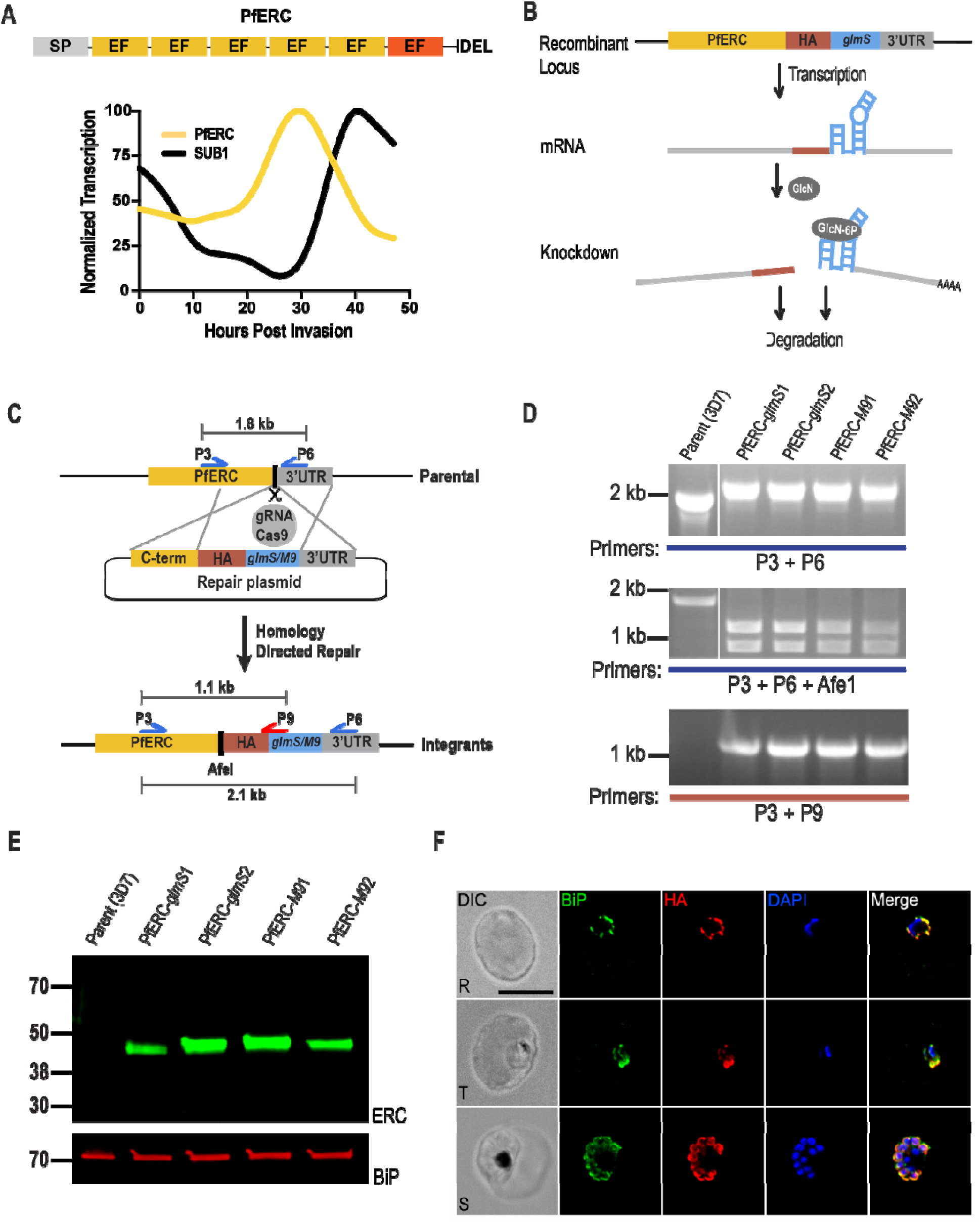
Generating PfERC-*glmS*/M9 mutant parasites. (A) Schematic representation of the domain structure of PfERC and its transcription profile. PfERC contains a signal peptide, 5 EF hands, and an ER-retention signal. Transcription data from *PlasmoDB* (26), were normalized to the highest expression value of the total abundance of the transcription. The expression profile of PfERC and an egress protease, SUB1, are shown. (B) Mechanism of the conditional expression of PfERC using the *glmS* ribozyme system. *glmS* is an inactive ribozyme that is transcribed, but not translated, with the mRNA of a protein of interest. Addition of glucosamine (GlcN) leads to its phosphorylation within the cell to glucosamine-6-phosphate (GlcN-6P). GlcN-6P binds to the transcribed PfERC-*glmS* mRNA and the *glmS* ribozyme is activated and cleaves itself from the mRNA. This leads to disassociation of the mRNA from its poly-A tail and leads to the degradation of target specific mRNA. The resulting decline in mRNA levels leads to reduced protein levels and, thus, loss of gene expression. As a control, we generated parasite lines containing a mutated version of the *glmS* ribozyme, called *M9*, which cannot cleave itself upon binding of GlcN. (C) Using the CRISPR/Cas9 system and a guide RNA targeting the PfERC gene, we induced a double stranded break in the PfERC locus that was repaired by a donor plasmid containing homology templates to the PfERC locus and appended a C-terminal 3XHA tag, the ER-retention signal (SDEL), and a stop codon followed by the *glmS* or *M9* sequence to the targeted gene. The location of diagnostic primers used to demonstrate the repair of the locus via double cross-over homologous integration are also shown (P3, P6 and P9). (D) PCR analysis of the generated mutants using specific primers (P3+P6; Table S1) in the C-terminus and 3’UTR of PfERC shows integration of the HA tag and *glms*/*M9* ribozymes into the PfERC locus. Modification of PfERC gene introduces an AfeI restriction enzyme site in this locus that is absent in the parental line. Digesting the PCR products (using AfeI) resulting from amplification using primers P3+P6 shows that AfeI is able to digest the PCR products from our mutants but not the parental line. PCR analysis using another primer pair (P3+P9) that sits on the *glmS*/*M9* sequence shows that amplification only occurs in the mutants but not in the parental line. (E) Western blot of lysates isolated from two independent clones and the parental line (3D7) probed with anti-HA antibodies show that the PfERC gene was tagged with HA in the mutants but not the parental line. PfBiP was the loading control. (F) Representative IFA of PfERC-*M9* parasites showing that tagged PfERC localizes to the ER as shown with co-localization with the ER chaperone BiP in all asexual stages of the parasite. From left to right, the images are phase-contrast, anti-BiP antibody (green), anti-HA antibody (red), and DAPI (blue), and fluorescence merge. Abbreviations: R, rings; T, trophozoites; S, schizonts. Scale bar, 5µm.

### Generating conditional mutants of PfERC

In order to determine the biological role of PfERC, we used CRISPR/Cas9 gene editing to generate conditional mutants of PfERC. In these parasite lines, the endogenous locus of PfERC was tagged with the inducible ribozyme, *glmS* or the inactive version of the ribozyme, *M9* (termed PfERC-*glmS* and PfERC-*M9* respectively) (Figure 1B and 1C) (27). PCR analysis of DNA isolated from PfERC-*glmS* and PfERC-*M9* parasite clones from two independent transfections demonstrate the correct insertion of the hemagglutinin (HA) tag and the *glmS/M9* ribozymes at the endogenous PfERC locus (Figure 1D). We detected expression of PfERC fused to the HA tag in the PfERC-*glmS* and PfERC-*M9* clones at the expected size and but not in the parental line (Figure 1E). Immunofluorescence microscopy confirmed that PfERC localized to the ER by co-staining with anti-HA and anti-BiP antibodies (Figure 1F).

To determine if PfERC was essential for intraerythrocytic survival, we grew asynchronous PfERC-*glmS* and PfERC-*M9* parasites in the presence of glucosamine (GlcN), which activates the *glmS* ribozyme leading to mRNA cleavage (Figure 1B). We observed a reproducible reduction of PfERC expression in PfERC-*glmS* parasites while there was no reduction in PfERC expression in PfERC-*M9* parasites grown under identical conditions (Figure 2A and 2B). Importantly, this reduction in PfERC levels inhibited the asexual expansion of PfERC-*glmS* parasites, while the PfERC-*M9* parasites were able to grow normally under the same conditions (Figure 2C). This inhibition of the asexual growth of PfERC-*glmS* parasites was dose-dependent upon GlcN (Supplementary figure 2A).

**Figure 2:**
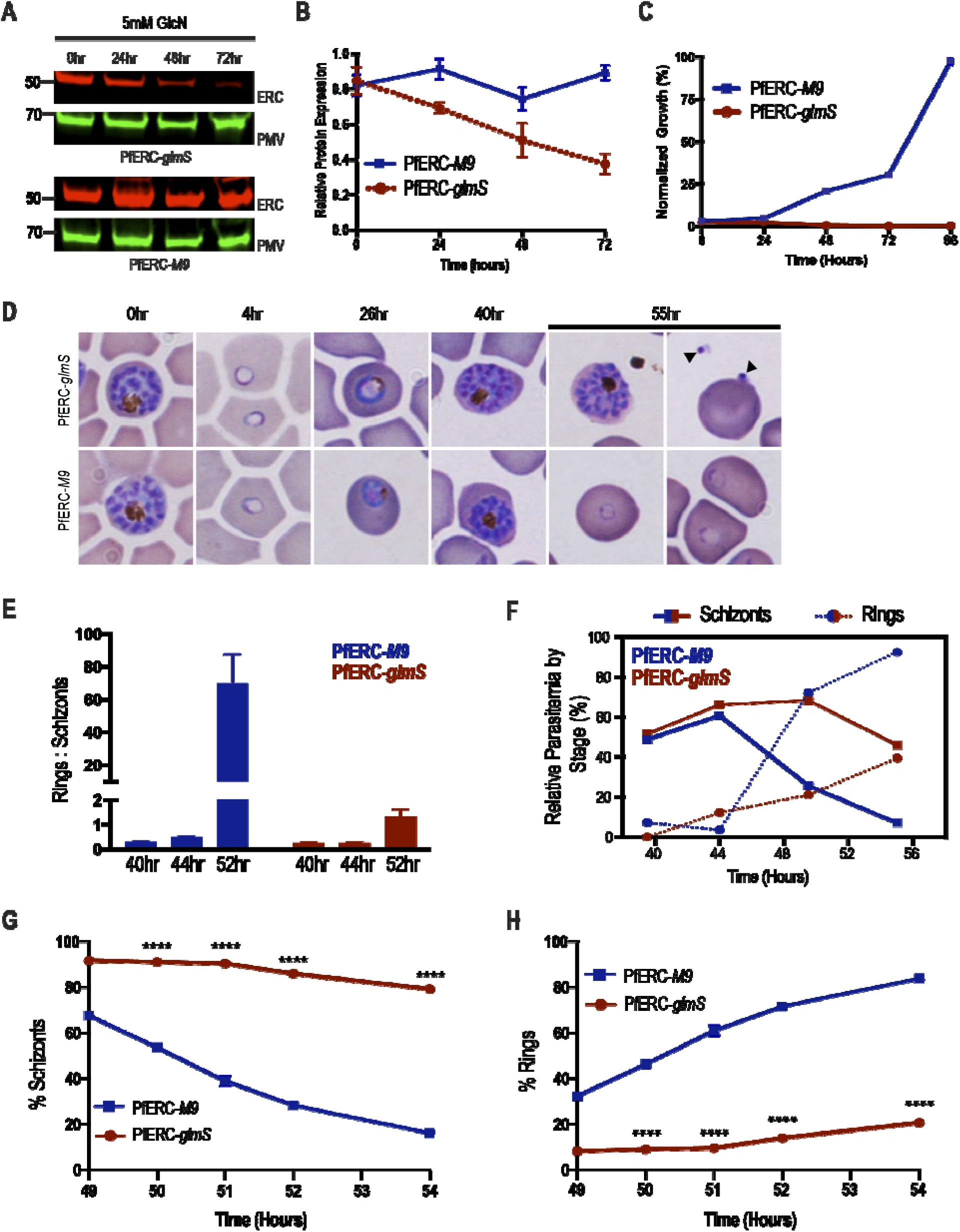
PfERC mutants fail to transition from schizonts to rings. (A) Western blot of parasite lysates isolated from PfERC-*glmS* and PfERC-*M9* parasites grown in the presence of 7.5 mM GlcN and probed with anti-HA antibodies (red) and anti-Plasmepsin V antibodies (PMV; green). One representative experiment out of four is shown. (B) Quantification of changes in expression of PfERC in PfERC-*glmS* and PfERC-*M9* parasites after addition of GlcN, as shown in (A). Data were normalized to the loading control (PMV) and shown as mean ± SEM (n=4 biological replicates). (C) Growth of asynchronous PfERC-*glmS* and PfERC-*M9* parasites incubated with 5mM GlcN, over 5 days, were observed using flow cytometry. Data are normalized to parasites grown without GlcN and are represented as the mean ± SEM (n=3). (D) Representative Hema-3 stained blood smears of synchronous PfERC-*glmS* and PfERC-*M9* parasites grown in the presence of GlcN (n=2 biological replicates). (E) GlcN was added to synchronous PfERC-*glmS* and PfERC-*M9* schizonts, and parasite stages were determined using flow cytometry. The ratio of rings to schizonts was calculated using the number of rings and schizonts observed at each time point. Data are represented as the mean ± SEM (n=3 biological replicates). (F) Hema-3 stained blood smears of synchronous PfERC-*glmS* and PfERC-*M9* parasites grown in the presence of GlcN (shown in D) were manually counted. The amount of each lifecycle stage (ring, trophozoite, and schizont) was determined as a percentage of the total number of parasites for each time point. (G, H) GlcN was added to synchronous PfERC-*glmS* and PfERC-*M9* schizonts, and parasite stages were determined using flow cytometry. At each time point, cells were fixed and stained with the DNA dye, Hoescht 33342, to distinguish between ring strage parasites (1N) and schizont stage parasites (16-32N). One representative experiment out of three are shown. Data are represented as the mean ± SEM (n=3 technical replicates; ****P<0.0001 2-way ANOVA).

### PfERC is essential for schizont to ring transition

Since our data show that PfERC was essential for growth within the host RBC, we used synchronous parasites to determine which asexual stage was affected by knockdown. We added GlcN to synchronized schizonts and observed the morphological development of the asexual stages at regular intervals during the intraerythrocytic life cycle (Figure 2D). All intracellular stages were morphologically normal in both PfERC-*glmS* and PfERC-*M9* parasites grown with GlcN (Figure 2D and Supplementary Figure 2B). However, 55hrs after addition of GlcN, the PfERC-*glmS* parasites remained either as morphologically normal schizonts or were observed as daughter merozoites in the extracellular space as well as some that were attached to RBCs (Figure 2D). On the other hand, PfERC-*M9* parasites were able to egress and re-invade fresh RBCs and developed into ring stage parasites (Figure 2D and Supplementary figure 2B).

These data suggest that knockdown of PfERC resulted in a defect in the conversion of schizonts into rings. To test this, we induced knockdown and observed the conversion of schizonts into rings via flow cytometry at 44, 48, and 56 h post-addition of GlcN. We found that over the course of 12 hours, PfERC-*M9* parasites transitioned from schizonts to rings as determined by the ring:schizont ratio while PfERC-*glmS* parasites were unable to convert from schizonts into rings resulting in a drastically reduced ratio (Figure 2E). Using synchronized PfERC-*glmS* and PfERC-*M9* parasites, treated as in Figure 2D, we observed the final hours of the asexual lifecycle using thin blood smears and quantified parasites using flow cytometry (Supplementary Figure 2B and Figure 2F-H). These data show that there was a delay in the disappearance of the morphologically normal PfERC-*glmS* schizonts over the final few hours of the asexual life cycle compared to PfERC-*M9* schizonts, suggesting that knockdown of PfERC led to a defect in egress (Figure 2F and 2G). Consequently, the delayed egress lead to reduced numbers of ring stage parasites in PfERC-*glmS* parasites unlike PfERC-*M9* parasites (Figure 2F and 2H).

### PfERC is not required for calcium storage

Since PfERC resides in the ER and possesses Ca^2+^ binding domains, we hypothesized that PfERC is required for egress because it plays a role in Ca^2+^ homeostasis in the ER. To test this model, synchronized PfERC-*glmS* and PfERC-*M9* schizonts were incubated with GlcN and allowed to proceed through one asexual cycle until they formed schizonts again. The second cycle schizonts were isolated using saponin lysis and loaded with Fluo-4AM to measure cytosolic Ca^2+^(Supplementary Figure 3A). To assess if the storage of Ca^2+^ in the ER of the parasite was affected by knockdown of PfERC, we added the SERCA-Ca^2+^ ATPase inhibitor, Cyclopiazonic acid (CPA), to these saponin-isolated parasites (Supplementary Figure 3A and 3B) (28). Inhibiting the SERCA-Ca^2+^ ATPase allows Ca^2+^ stored in the ER to leak into the cytoplasm, which results in a detectable change in the fluorescence of Fluo-4AM (Supplementary Figure 3B). Our measurements show that there was no difference in the amount of Ca^2+^ that leaked from the parasite ER, upon SERCA-Ca^2+^ ATPase inhibition, between PfERC-*glmS* and PfERC-*M9* schizonts (Supplementary Figure 3B).

To test if there was a defect in Ca^2+^ storage in neutral stores, we used the ionophore, Ionomycin, which releases Ca^2+^ from all neutral stores in the cell and measured the release of Ca^2+^ into the cytoplasm of PfERC-*glmS* and PfERC-*M9* schizonts. The parasites were isolated as described above and the changes in cytoplasmic Ca^2+^ were measured using Fluo-4AM (Supplementary Figure 3A and 3C). Again, we did not observe any difference in the amount of Ca^2+^ released into the cytoplasm of PfERC-*glmS* and PfERC-*M9* schizonts treated with ionomycin (Supplementary Figure 3C). These data suggest that the availability of free Ca^2+^ in the ER (or other neutral Ca^2+^ stores) of *P. falciparum* is not affected by knockdown of PfERC. Furthermore, these data suggest that the observed egress defect upon PfERC knockdown was not a result of disequilibrium of Ca^2+^in the parasite ER.

### PfERC is required for PVM breakdown

Since we could not observe a defect in ER Ca^2+^ storage upon knockdown, we further analyzed how egress of PfERC-*glmS* parasites was failing during knockdown. Egress of daughter merozoites from the infected RBC is an ordered and rapid process where the PVM breakdown precedes the disruption of RBCM (Figure 3A) (11). Therefore, we took synchronized PfERC-*glmS* and PfERC-*M9* schizonts and initiated knockdown with addition of GlcN. These schizonts were allowed to reinvade fresh RBCs and proceed through the asexual stages for 48 hours until they developed into schizonts again. Then, these second cycle schizonts were incubated with inhibitors that block key steps during egress of *P. falciparum* (Figure 3A). To ensure synchronized egress, we used reversible inhibitors of PKG, Compound 1 (C1) or Compound 2 (C2), because inhibition of PKG allows merozoites to develop normally but prevents them from initiating egress (Figure 3A) (6, 11). We used flow cytometry to observe PfERC-*glmS* and PfERC-*M9* schizonts after washing off C1 and saw that there was a delay in the egress of PfERC-*glmS* schizonts while the majority (>60%) of the PfERC-*M9* schizonts were able to complete egress within two hours after washout of C1 (Figure 3B). Consequenctly, this led to a loss of ring formation in PfERC-*glmS* parasites as compared to PfERC-*M9* parasites (Supplementary Figure 4A). Removal of C1 initiates the breakdown of the PVM followed by RBCM rupture (Figure 3A), suggesting that PfERC-*glmS* parasites fail to breach one of these membranes down despite removal of the PKG inhibitor.

**Figure 3:**
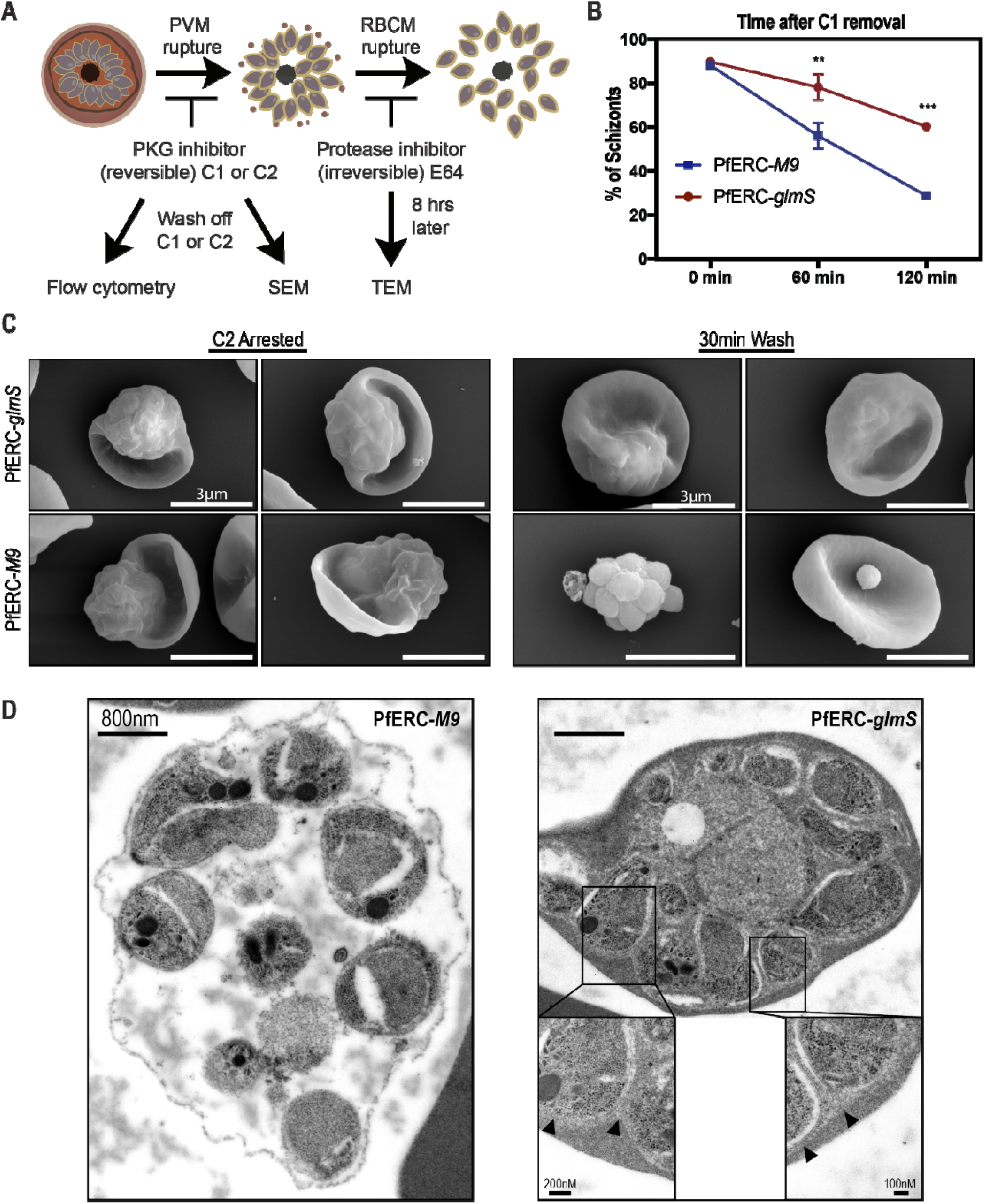
PfERC knockdown inhibits PVM breakdown. (A) Schematic showing the experimental layout to study the effect of PfERC knockdown using specific compounds that inhibit egress in parasites. Abbreviations: C1/2-PKG inhibitors, compound 1 or compound 2; SEM-Scanning Electron Microscopy; TEM-Transmission Electron Microscopy. (B) As shown in (A), synchronized PfERC-*glmS* and PfERC-*M9* schizonts were grown in the presence of GlcN and second cycle schizonts were observed by flow cytometry after removal of C1 (time 0hr). Schizonts were quantified as a percentage of the total amount of parasites as determined by flow cytometry. Data are represented as the mean ± SEM (n=3 biological replicates; **P<0.01, ***P<0.001 2-way ANOVA). (C) Representative SEM images of C2 arrested PfERC-*glmS* (n=4 biological replicates) and PfERC-*M9* (n=4 biological replicates) mutants fixed immediately after washing off C2 and after 30mins, as shown in (A). (D) Representative TEM images of PfERC-*glmS* (n=2 biological replicates) and PfERC-*M9* (n=2 biological replicates) schizonts incubated with E-64, as shown in (A). Scale bar, 800nm. Higher magnification images for PfERC-*glmS* parasites where the PVM is marked by black arrowheads are shown in the inset.

Therefore, we tested whether PfERC knockdown prevented rupture of PVM or if PfERC was required for RBCM breakdown (Figure 3A). PfERC-*glmS* and PfERC-*M9* schizonts (where knockdown had been initiated in the previous cycle) were incubated with C2 (6, 11) and observed by scanning electron microscopy (SEM) (Figure 3A and 3C). We observed that parasites treated with C2 were morphologically identical and had developed into mature schizonts within the PVM inside the RBC (Figure 3C). Then, we washed C2 from the parasites and observed these schizonts after 30 mins by SEM (Figure 3C). During this time period, the majority of PfERC-*M9* schizonts were able to initiate egress after removal of C2 and we observed free merozoites attached to the RBC as well as clusters of merozoites that had broken out of the PVM but were contained by a collapsed RBCM wrapped around them (Figure 3C and Supplementary Figure 4B). In contrast, the majority of PfERC-*glmS* schizonts were still stuck within the RBC and looked identical to the C2 arrested schizonts, suggesting that they had not initiated egress even though PKG was no longer inhibited (Figure 3C and Supplementary Figure 4). These data suggest that knockdown of PfERC blocks egress at an early step, perhaps blocking the rupture of the PVM (Figure 3C).

We directly observed if breakdown of the PVM was impacted by knockdown of PfERC using transmission electron microscopy (TEM) (Figure 3D). Knockdown was induced by adding GlcN to PfERC-*glmS* and PfERC-*M9* schizonts and these parasites were allowed to go through one asexual cycle and develop into schizonts again 48hrs later. These schizonts were prevented from completing egress using the irreversible cysteine protease inhibitor, E-64 (Figure 3A). This inhibitor blocks the breakdown of the RBCM but allows both the breakdown of PVM and poration of the RBCM, which results in the loss of the electron dense contents of the infected RBC (Figure 3A) (11, 29, 30). Our results show that the PfERC-*M9* schizonts were able to break down the PVM as well as proceed with the poration of the RBCM after an 8-hour incubation with E-64, while the PfERC-*glmS* mutants were unable to proceed through the first step of egress and failed to rupture the PVM (Figure 3D). Overall, these data demonstrate that PfERC function is critical for the breakdown of the PVM (Figure 3C and 3D).

### SUB1 maturation requires PfERC

Electron microscopy data show that knockdown of PfERC prevents the breakdown of the PVM (Figure 3). A key event required for PVM breakdown is the proteolytic processing of SUB1, which is required to start a proteolytic cascade that ends in the release of merozoites from the infected RBC (14, 29). Therefore, we tested if knockdown of PfERC affects processing of PfSUB1. This protease is processed twice, once in the ER, where it undergoes a Ca^2+^-dependent autocatalytic processing from its zymogen form (83-kDa) into a 54-kDa semi-proenzyme form (p54) (18, 19, 31). From the ER, SUB1 is transported to the egress-related secretory vesicles, the exonemes, which are secreted into the PV to initiate breakdown of the PV membrane. It is proposed that during trafficking of SUB1 to exonemes, it is processed by PMX from its semi-proenzyme form (p54) to its mature form (p47) (16, 17). The secretion of mature p47 form of SUB1 initiates the breakdown of the PVM (14, 31). Given that one CREC family member has been shown to transiently interact with a subtilisin like protease in mammalian cells (25), we hypothesized that PfERC is required for one of the proteolytic maturation steps of SUB1, most likely the first Ca^2+^-dependent autocatalytic processing step in the ER.

To test this hypothesis, PfERC-*glmS* and PfERC-*M9* schizonts were incubated with GlcN and allowed to progress through one asexual growth cycle (48 hours) to develop into schizonts again. Lysates from these synchronized schizonts were separated on a Western blot and probed with antibodies against SUB1 (Figure 4A and Supplementary Figure 5A). No change was observed in the Ca^2+^-dependent autoprocessing of the zymogen-form of SUB1 into the semi-proenzyme (p54) form (Figure 5A and Supplementary Figure 5A). Surprisingly, we observed a reproducible and significant decrease in the processing of SUB1 from p54 to the p47 form in PfERC-*glmS* parasites (Figure 4A and 4B). Compared to PfERC-*M9* parasites, there was a >50% decrease in the amount of processed SUB1 (p47) in PfERC-*glmS* parasites (Figure 4B). This effect was also observed in cells treated with Compound 1, suggesting that PfERC is required for SUB1 processing prior to secretion of exonemes (Figure 5C and 5D). Taken together, our data suggest that PfERC is essential for the proteolytic maturation of SUB1.

**Figure 4:**
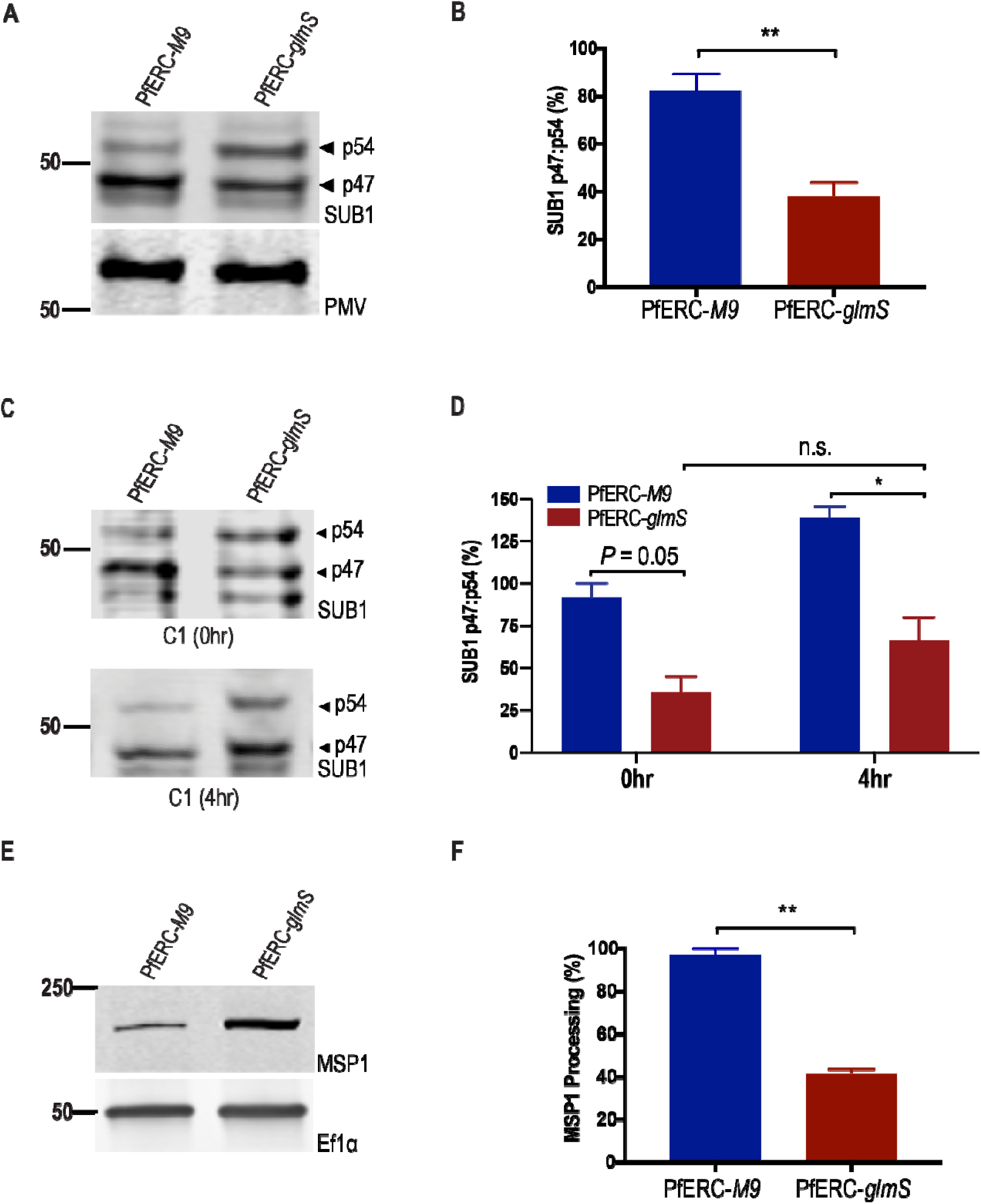
PfERC knockdown inhibits SUB1 and MSP1 processing. (A) Western blot of parasite lysates isolated from PfERC-*glmS* and PfERC-*M9* schizonts grown in the presence of GlcN for 48 hours and probed with anti-SUB1 antibodies (top panel) and anti-PMV (loading control, bottom panel). One representative experiment out of four is shown. The protein marker sizes that co-migrated with the probed protein are shown on the left. (B) Quantification of SUB1 processing in PfERC-*glmS* and PfERC-*M9* parasites over time after addition of GlcN, as shown in (A). Data were normalized to the ratio of processed SUB1 (p47:p54) of PfERC-*M9* parasites and are represented as mean ± SEM (n=4 biological replicates; **P<0.005 unpaired t-test). (C) Western blot of parasite lysates isolated from PfERC-*glmS* and PfERC-*M9* schizonts grown in the presence of GlcN for 48hrs and then incubated with Compound 1. Samples were taken either 0hrs or 4hrs post addition of Compound 1. (D) Quantification of SUB1 processing in PfERC-*glmS* and PfERC-*M9* parasites incubated with Compound 1 as shown in (C). Data were normalized to the ratio of processed SUB1 (p47:p54) of PfERC-*M9* parasites at 0hr and represented as ± SEM. (n=2 biological replicates; \**P*<0.05 one-way ANOVA). (E) Western blot of parasite lysates isolated from PfERC-*glmS* and PfERC-*M9* schizonts grown in the presence of GlcN for 48 hours and probed with anti-MSP1 12.4 antibodies (top panel) and with anti-EF1α (loading control, bottom panel). One representative experiment out of two is shown. (F) Quantification of unprocessed (or full length) MSP1 in PfERC-*glmS* and PfERC-*M9* parasites after addition of GlcN, as shown in (E). Data were normalized to the loading control (EF1α) and are represented as mean ± SEM (n=2 biological replicates; **P<0.005 unpaired *t*-test).

**Figure 5:**
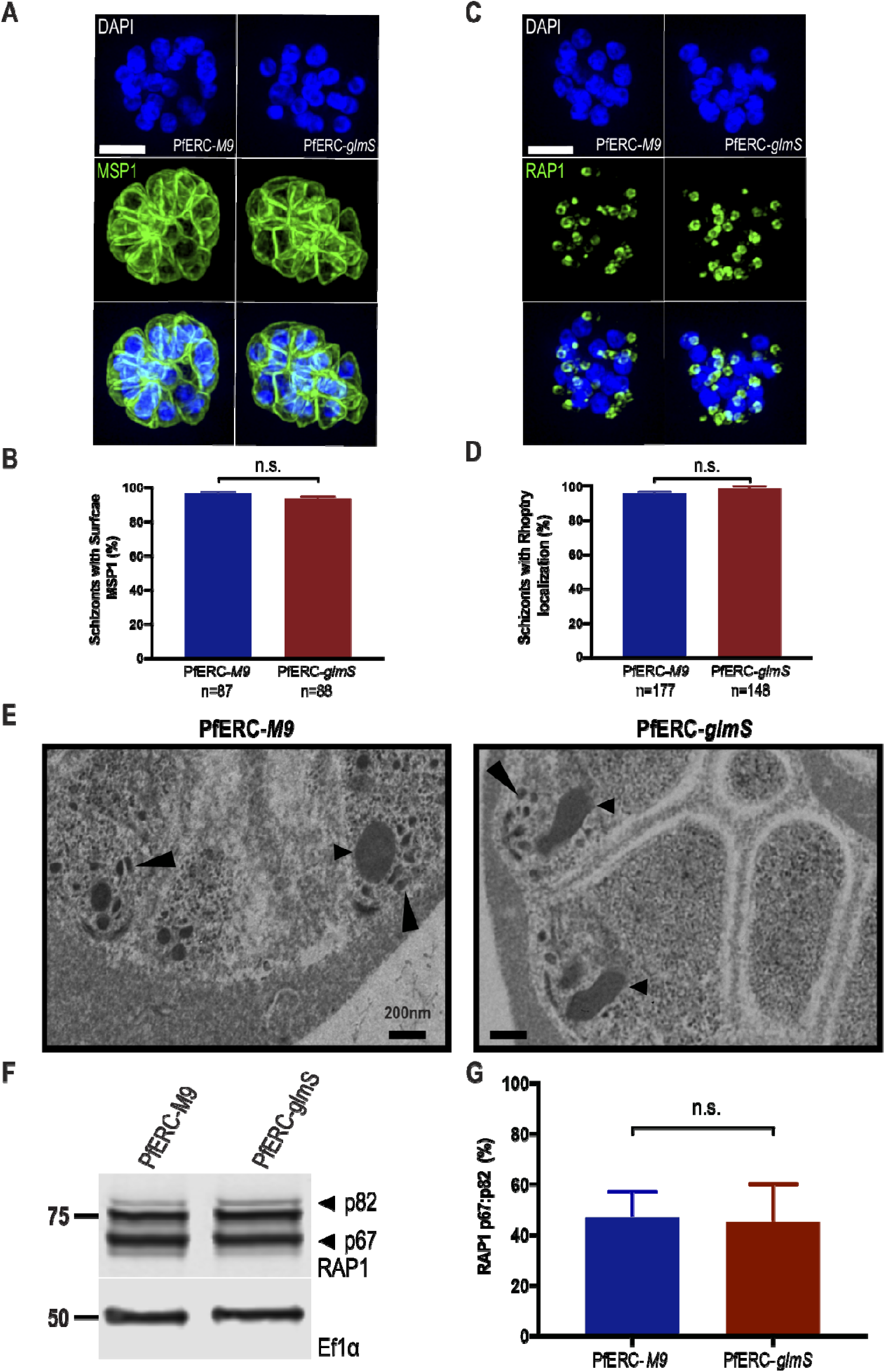
PfERC is not required for protein trafficking or organellar biogenesis. (A) Representative Super-Resolution SIM images of PfERC-*glmS* and PfERC-*M9* schizonts incubated with GlcN for 48hrs and stained with anti-MSP1 and the nuclear stain, DAPI. From top to bottom, the images are anti-MSP1 (green), DAPI (blue), and fluorescence merge. Scale bar 2µm. (B) The surface localization of MSP1 was quantified in PfERC-*glmS* and PfERC-*M9* schizonts incubated with GlcN for 48hrs and stained as shown in (A). Data are represented as mean ± SEM (n=2 biological replicates; n.s.=non-significant, unpaired *t*-test). (C) Representative Super-Resolution SIM images of PfERC-*glmS* and PfERC-*M9* schizonts incubated with GlcN for 48hrs and stained with anti-RAP1 and the nuclear stain, DAPI. From top to bottom, the images are anti-RAP1 (green), DAPI (blue), and fluorescence merge. Scale bar 2µm. (D) The rhoptry localization of RAP1 was quantified in PfERC-*glmS* and PfERC-*M9* schizonts incubated with GlcN for 48hrs and stained as shown in (C). Data are represented as mean ± SEM (n=3 biological replicates; n.s.=non-significant, unpaired *t*-test). (E) Representative TEM images of synchronized PfERC-*glmS* and PfERC-*M9* schizonts grown for 48hrs with GlcN and incubated with C1 for 4 hours, as shown in Figure 3A (n=2 biological replicates). Small arrowheads point to rhoptries, large arrowheads to micronemes. Scale bar, 200nm. (F) Western blot of parasite lysates isolated from PfERC-*glmS* and PfERC-*M9* schizonts grown in the presence of GlcN for 48 hours and probed with anti-RAP1 (top panel) and anti-EF1α (loading control, bottom panel). One representative experiment out of five is shown. The protein marker sizes that co-migrated with the probed protein are shown on the left. (G) Quantification of RAP1 processing in PfERC-*glmS* and PfERC-*M9* parasites incubated with GlcN, as shown in (G). Data were normalized to the ratio of processed RAP1 (p67:p82) of PfERC-*M9* parasites and are represented as mean ± SEM (n=5 biological replicates; n.s=non-significant, unpaired t-test).

Since we observed the presence of some mature SUB1 in PfERC-*glmS* parasites (Figure 4A), we tested if the activity of SUB1 was affected upon knockdown of PfERC by assaying for the processing of a known SUB1 substrate, the merozoite surface protein 1 (MSP1). MSP1 is required for the initial attachment of merozoites onto RBCs and it has been shown that correct processing of MSP1 by SUB1, is also required for efficient egress as it plays a role in breakdown of the RBC cytoskeleton after release from the PVM (32–35). Lysates from synchronized second-cycle PfERC-*glmS* and PfERC-*M9* schizonts, treated as above, were separated on a Western blot and probed using anti-MSP1 antibodies (Figure 4E and Supplementary Figure 5B and 5C). Our data show that there was significant inhibition of MSP1 processing in PfERC-*glmS* parasites as compared to PfERC-*M9* parasites after knockdown (Figure 4F and Supplementary Figure 5B and 5C). These data reveal that knockdown of PfERC leads to defects in SUB1 processing and activity, and consequently, MSP1 processing (Figure 4 and Supplementary Figure 5B and 5C).

### PfERC is not required for protein trafficking or organelle biogenesis

MSP1 is a GPI-anchored merozoite membrane protein that is presumably processed by SUB1 once the protease is secreted into the PV (29, 32). Therefore, we wanted to verify that knockdown of PfERC led to a specific defect in the egress cascade and is not due to a block in protein trafficking via the ER or defects in the biogenesis of egress and invasion related organelles. To address this, we used super resolution structured illumination microscopy (SR-SIM) to observe if there was a difference in the surface expression of MSP1 between PfERC-*glmS* and PfERC-*M9* schizonts upon knockdown of PfERC (Figure 5A). As before, knockdown was initiated in synchronized PfERC-*glmS* and PfERC-*M9* schizonts and after 48 hours, these schizonts were stained with anti-MSP1 antibodies. Our data shows that there was no difference in the trafficking of MSP1 to the surface of developing PfERC-*glmS* or PfERC-*M9* merozoites after knockdown (Figure 5A and 5B). Further, the localization of the rhoptry protein, RAP1, which also traffics through the ER, was observed in these schizonts using SR-SIM. Our data show that there was no difference in the localization of RAP1 in schizonts between PfERC-*glmS* and PfERC-*M9* parasites suggesting that the knockdown of PfERC does not cause a generalized defect in the secretory pathway (Figure 5C and 5D).

As the ER produces the lipid membranes required for generating organelles essential for egress and invasion, we observed if organelle biogenesis was inhibited upon PfERC knockdown. To test this, knockdown was initiated in synchronized PfERC-*glmS* and PfERC-*M9* schizonts and after 48 hours, these schizonts were treated with Compound 1 for 4hrs. Then, we observed these C1-treated schizonts using TEM. In these PfERC-*glmS* and PfERC-*M9* parasites both micronemes and rhoptries remain morphologically intact (Figure 5E). Likewise, we observed morphologically intact micronemes and rhoptries in PfERC-*glmS* and PfERC-*M9* schizonts that were further incubated with E-64 for 8 hours (Supplementary Figure 6).

Even though we did not observe any morphological defects in the formation of organelles it is possible that rhoptry proteins may not be processed correctly upon PfERC knockdown. For example, after transport to the rhoptry, the rhoptry-bulb protein, RAP1, undergoes essential proteolytic cleavage by the aspartic protease, Plasmepsin IX (PMIX), from a pro-form (p83) to a mature form (p67) (16, 17, 36, 37). Therefore, we tested if RAP1 processing was inhibited by knockdown of PfERC using Western blot analysis (Figure 5F and Supplementary Figure 5D). Our data show that the proteolytic processing of RAP1 was not inhibited by the knockdown of PfERC (Figure 5G), showing that knockdown does not lead to a generalized defect in the processing of all proteins that traverse through the secretory pathway. Together, these data suggest that the knockdown of PfERC does not lead to defects in organelle biogenesis (Figure 3D, Figure 5E, and Supplementary Figure 6).

### PfERC is required for AMA1 processing but not secretion

The invasion ligand, AMA1, is essential for invasion-competent merozoites because processed AMA1 is required for the formation of the tight junction between the parasite and the RBC (38, 39). It is trafficked from micronemes to the merozoite surface prior to egress, where it is processed from its pro-form (p83) to its mature form (p66) by an unknown protease (40–42). Studies have shown that secretion of micronemes require Ca^2+^ signaling pathways (9) and, although our data suggest that PfERC is not required for Ca^2+^ storage in the ER, we could not rule out a role for PfERC in Ca^2+^ signaling. Since the translocation of AMA1 is dependent upon this Ca^2+^ signaling pathway, we observed if AMA1 exocytosis was inhibited upon PfERC knockdown (6, 12). Synchronized PfERC-*M9* or PfERC-*glmS* schizonts where knockdown had been initiated the previous cycle were incubated with C1 to achieve tight synchronization. Then, C1 was washed off and the parasites were incubated with E-64 to trap merozoites that had initiated egress within the RBC membrane (12). Using immunofluorescence microscopy, we observed AMA1 localization in the either micronemes or on the surface of merozoites (Figure 7A). Our data show that there was no difference in the localization of AMA1 between PfERC-M9 or PfERC-glmS parasites, suggesting that PfERC is not required for the signaling necessary for vesicle secretion (Figure 6A and 6B).

**Figure 6:**
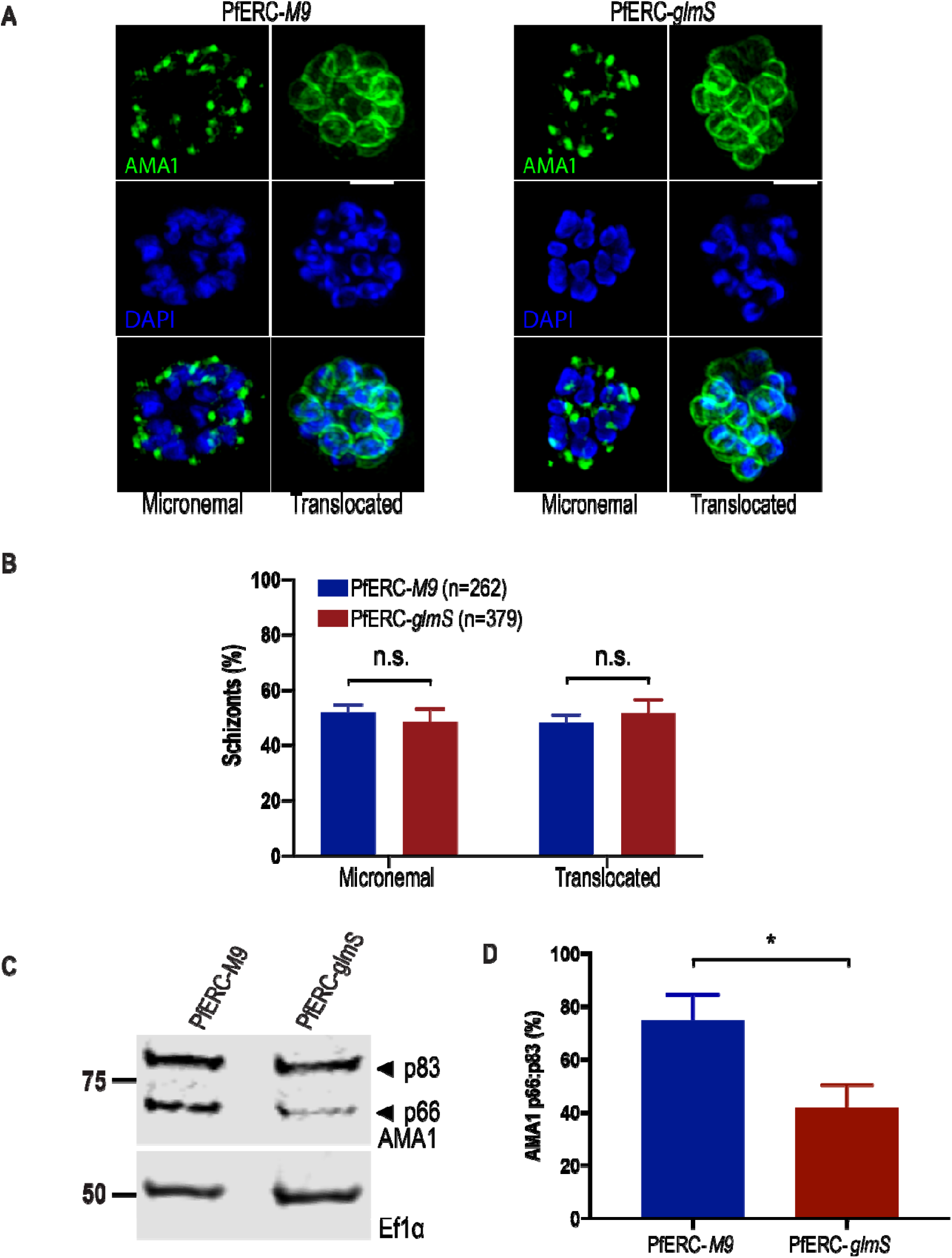
Proteolytic processing of AMA1 requires PfERC. (A) Parasites were grown in the presence of GlcN for 48hrs and then incubated with Compound 1 for 4hrs. Compound 1 was then removed and parasites were incubated further with E-64 for 6hrs and stained with anti-AMA1 as well as the nuclear stain, DAPI. Representative Super-Resolution SIM images of these PfERC-*glmS* and PfERC-*M9* schizonts are shown. From top to bottom, the images are anti-AMA1 (green), DAPI (blue), and fluorescence merge. Scale bar 2µm. (B) The micronemal and surface (or translocated) localization of AMA1 was quantified in PfERC-*glmS* and PfERC-*M9* schizonts as shown in (A). Data are represented as mean ± SEM (n=4 biological replicates; n.s.=non-significant, one-way ANOVA). (C) Western blot of parasite lysates isolated from PfERC-*glmS* and PfERC-*M9* schizonts grown in the presence of GlcN for 48 hours and probed with anti-AMA1 antibodies (top panel) and anti-EF1α antibodies (loading control, bottom panel). One representative experiment out of eight is shown. The protein marker sizes that co-migrated with the probed protein are shown on the left. (D) Quantification of AMA1 maturation in PfERC-*glmS* and PfERC-*M9* parasites incubated with GlcN, as shown in (C). Data were normalized to the ratio of processed AMA1 (p66:p83) in PfERC-*M9* parasites and are represented as mean ± SEM (n=8 biological replicates; *P<0.05 unpaired t-test).

**Figure 7:**
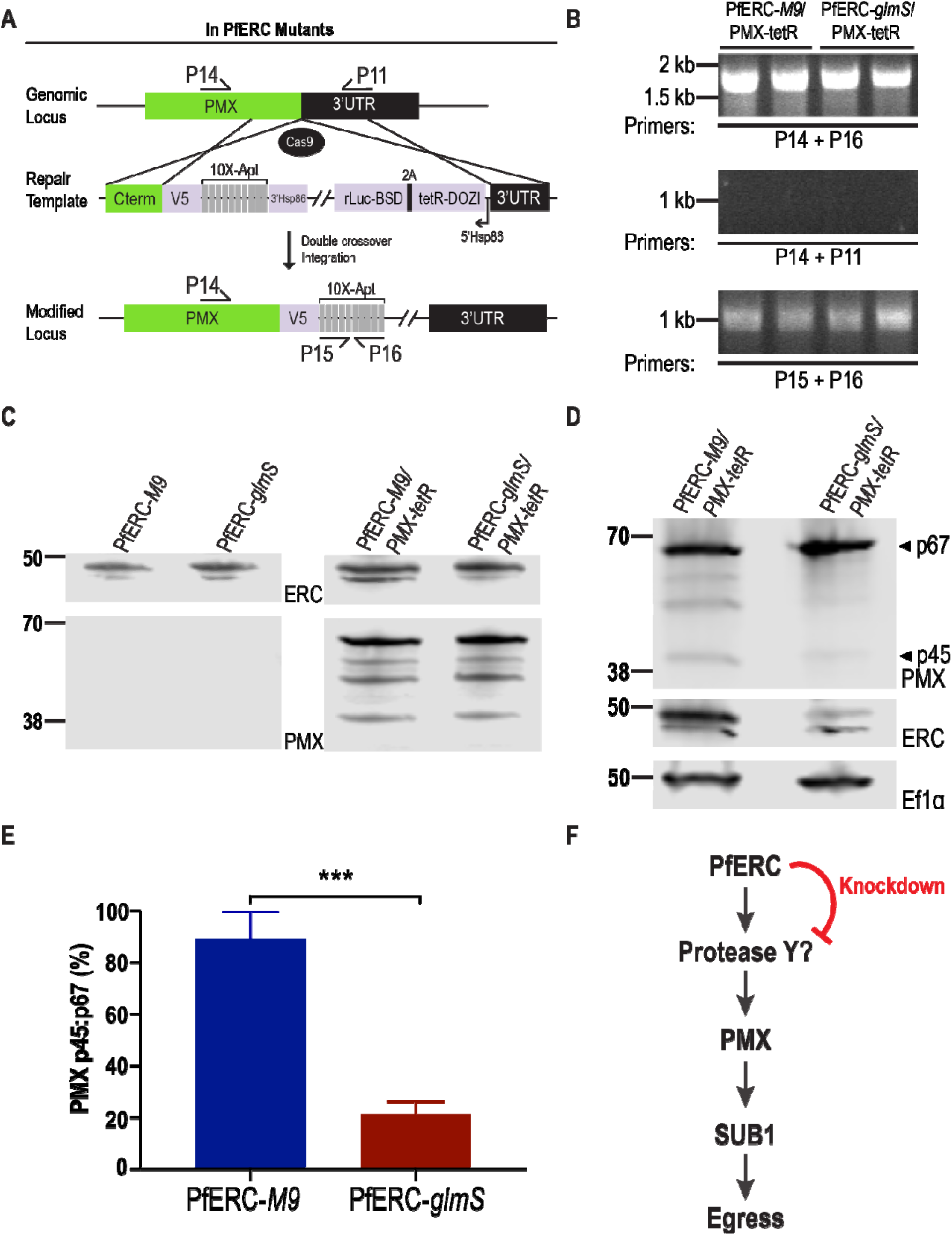
PfERC is required for PMX cleavage. A) Using the CRISPR/Cas9 system and a guide RNA targeting the PMX gene, we induced a double stranded break in the PMX locus that was repaired by a donor plasmid containing homology templates to the PMX locus and appended a C-terminal 3XV5 tag and a stop codon followed by the 10X aptamer sequences. The location of diagnostic primers used to demonstrate the repair of the locus via double cross-over homologous integration are also shown (P11, P14, P15 and P16). (B) PCR analysis of the generated mutants (2 clones each from the PfERC-*glmS* background and the PfERC-*M9* background) using specific primers (P14+P11; Table S1) in the C-terminus and aptamer region show integration of the plasmid into the PMX locus. PCR analysis using specific primers in the C-terminus and 3’UTR of PMX show the absence of wildtype parasites in the clonal population and PCR analysis specific to the aptamer region (P15+P16) show correct number of aptamers in our clones. (C) Representative Western blot of lysates isolated from the parental PfERC-*glmS* and PfERC-*M9* clones as well as the PfERC-*glmS*/PMX-tetR and PfERC-*M9*/PMX-tetR double mutants probed with anti-V5 antibodies show that the PfPMX gene was tagged with V5 in the double mutants but not the parental line. D) Western blot of lysates isolated from PfERC-*glmS*/PMX-tetR and PfERC-*M9*/PMX-tetR schizonts grown in the presence of GlcN for 48 hours and probed with anti-V5 (top panel), anti-HA (middle panel), and anti-Ef1α (loading control, bottom panel) antibodies. One representative experiment out of four is shown. E) Quantification of PMX processing in PfERC-*glmS*/PMX-tetR and PfERC-*M9*/PMX-tetR parasites 48 hours after addition of GlcN, as shown in (D). Data were normalized to the ratio of processed PMX (p45:p67) of PfERC-*M9*/PMX-tetR parasites and are represented as mean ± SEM (n=4 biological replicates; ***P<0.001 unpaired t-test). F) A proposed model for the role of PfERC in egress. PfERC regulates the activity an unknown protease (Protease Y) that cleaves PMX. PMX then cleaves SUB1, eventually leading to egress of the parasite. Knockdown of PfERC prevents the activity of Protease Y and thus inhibiting the proteolytic cascade required for egress.

The proteolytic processing of AMA1 is critical for its function during invasion (43). Therefore, we tested if the processing of the AMA1 was affected upon knockdown of PfERC. As before, after initiating knockdown in synchronized schizonts, we isolated lysates from second cycle PfERC-*glmS* and PfERC-*M9* schizonts on a Western blot and probed it with anti-AMA1 antibodies (Figure 6C and Supplementary Figure 5E). We observed a significant reduction in the proteolytic processing of AMA1 upon knockdown (Figure 6C). Indeed, there was a >40% decrease in the processing of AMA1 in PfERC-*glmS* mutants compared to the PfERC-*M9* control (Figure 6D). These data suggest that PfERC is required for the proteolytic maturation of AMA1 during egress.

### PfERC knockdown inhibits PMX maturation

Our data show that PfERC knockdown inhibits SUB1 maturation as well as processing of downstream substrates. Since SUB1 binds Ca^2+^ and this is required for its activity, one possibility is that PfERC directly regulates the proteolysis of SUB1. In mammalian cells, CREC family members have been shown to interact transiently with a subtilisin-like protease (25). Therefore, to test if PfERC interacts directly with SUB1, we performed co-IP experiments but failed to detect any interaction between these two proteins (Supplementary Figure 7).

This suggested that either the interaction is too transient to detect by co-IP or that PfERC works upstream of SUB1 via the recently discovered aspartic protease, PMX (16, 17). To test this, we generated double conditional mutant parasites where we tagged PMX with *tetR* aptamers in both PfERC-*glmS* and PfERC-*M9* parasites (termed PfERC-*glmS*/PMX-*tetR* and PfERC-*M9*/PMX-*tetR*) (Figure 7A) (44). This aspartic protease undergoes proteolytic maturation from 67kDa zymogen to a 45kDa active protease (16, 17). Our data show that we were successful in tagging PMX with the V5 tag in PfERC-*glmS*/PMX-*tetR* and PfERC-*M9*/PMX-*tetR* parasites (Figure 7B and Figure 7C). As before, after initiating PfERC knockdown in synchronized schizonts, we isolated lysates from second cycle PfERC-*glmS*/PMX-*tetR* and PfERC-*M9*/PMX-*tetR* schizonts and monitored PMX cleavage on a Western blot. We observed a reproducible and significant decrease in the processing of PMX from p67 to the p45 form in PfERC-*glmS*/PMX-*tetR* parasites (Figure 7D and 7E and Supplementary Figure 5F). No such deficiencies in PMX cleavage was observed in PfERC-*M9*/PMX-*tetR* parasites (Figure 7D and Figure 7E). Together, these data show that PfERC is essential for the proteolytic maturation of PMX during egress, placing it as the earliest known regulator in the egress proteolytic cascade in malaria parasites.

## Discussion

In this study, we revealed the biological role of a conserved Ca^2+^-binding protein that resides in the lumen of the ER of *Plasmodium falciparum*. Our data show that PfERC is essential for asexual replication of malaria parasites. Knockdown of PfERC did not affect the ring and trophozoite development but clearly inhibited the subsequent schizont-to-ring transition. Specifically, these data show that PfERC is required for egress from infected RBCs and potentially for invasion into host erythrocytes. This is consistent with data that suggest PfERC may be transcriptionally controlled by the invasion-specific transcription factor PfAP2-I (45). Knockdown of PfERC leads to defects in the processing of proteins critical for invasion of merozoites into the host RBC, namely, MSP1 and AMA1. However, any invasion defect is likely a secondary effect because several proteins critical for invasion are processed during egress (16, 17, 29, 32). Given the kinetic limitations of the conditional knockdown system, we cannot tease out a specific role for PfERC in invasion. As invasion occurs rapidly (<2mins), a potential specific invasion-related function of PfERC could be tested using a small molecule that specifically targets PfERC (46). Overall, these data show that PfERC is essential for egress of merozoites from the infected RBC.

During the formation of daughter merozoites in schizonts, several key egress and invasion related organelles essential for propagation of the infection are generated. The ER is thought to play a key role in the biogenesis of these organelles and the ER is responsible for transporting the essential proteins to these organelles (20, 21). As an ER-resident protein, knockdown of PfERC could affect several ER functions such as protein trafficking, organellar biogenesis, and Ca^2+^ signaling. Therefore, we tested if PfERC functions in the trafficking of proteins required for schizont to ring transition such as MSP1, AMA1, and RAP1. A defect in the secretory pathway would explain the observed deficiencies in the proteolytic processing of SUB1, MSP1 and AMA1, as transport out of the ER is required for their maturation (6, 19, 37). However, super-resolution and electron microscopy experiments show that proteins on the merozoite surface, micronemes, and rhoptries are trafficked normally and biogenesis of egress and invasion organelles is normal. Likewise, Western blot analysis showed that the proteolytic processing of a rhoptry protein, RAP1, which is processed after transport to the organelle, occurs normally upon knockdown of PfERC (16, 17). These data show that knockdown of PfERC does not result in a generalized defect in protein trafficking via the ER, or in organelle biogenesis.

CREC family members in the ER are known to regulate the function of Ca^2+^ pumps and channels such as the Ryanodine and IP_3_ receptors (47, 48). Therefore, one interesting possibility we considered was that PfERC may play a role in the signal-dependent release of Ca^2+^ from the ER. This is difficult to test in *Plasmodium* since there are no clear orthologs for a ligand-dependent Ca^2+^ channel in its genome (49). Intracellular Ca^2+^ stores are required for egress and invasion of malaria parasites since cell permeable Ca^2+^ chelators block egress of *Plasmodium* parasites from host RBCs (13, 50–53). Further, Ca^2+^ binding proteins in the parasite cytoplasm are essential for egress of malaria parasites, for example, the Ca^2+^ dependent protein kinase, PfCDPK5, is required for secretion of egress specific organelles, such as those containing AMA1 (12, 54). As PfCDPK5 is thought to be activated upon the signal-dependent release of intracellular Ca^2+^ into the cytoplasm (12), we tested if PfERC was required for exocytosis of AMA1-containing vesicles. The data suggest that PfERC is not required for the PfCDPK5-dependent translocation of AMA1 onto merozoite membrane. However, PfERC is required for the essential proteolytic maturation of AMA1, suggesting that this CREC family member regulates (directly or indirectly) the unknown protease that cleaves AMA1.

An essential enzyme vital for initiating egress is the protease SUB1 as this serine protease is required for the rupture of both the PVM and the RBCM (14, 29). SUB1 is produced as an 82-kDa zymogen in the ER, where it rapidly self-processes into a 54-kDa semi-proenzyme in the ER (31). If PfERC was needed for this autoprocessing event, then this would explain the observed knockdown phenotypes (29). Instead, our data show that PfERC was essential for the second processing step of SUB1, which produces the mature, active form of the protease (p54 to p47). This processing event occurs once is trafficked out of the ER suggesting a role for PfERC in SUB1 maturation once it leaves the ER (6, 13, 17, 18). The release of the mature SUB1 into the PV kickstarts the egress cascade (29) and the cGMP signaling pathway is thought to be essential for vesicle exocytosis via the release of intracellular Ca^2+^ stores (52).

Therefore, we tested whether PfERC plays a role in the cGMP signaling pathway, using the PKG inhibitor Compound 1, and show that PfERC knockdown inhibited SUB1 maturation even when PKG activity was inhibited. This data together with the experiments testing AMA1 translocation onto the merozoite membrane suggest that PfERC does not play a role in the Ca^2+^-dependent exocytosis of egress-specific organelles. Instead, our data suggest a model where PfERC plays a specific role in the maturation of SUB1 prior to its secretion into the PV.

One possibility is that PfERC is required for the maturation of a protease in this pathway that works upstream of SUB1. The recently discovered aspartic protease, PMX, cleaves SUB1 from p54 to p47 (16, 17). In turn, PMX itself is proteolytically processed from a 67kDa zymogen to a 45kDa active protease (16, 17). But unlike most aspartyl proteases, PMX does not autoprocess because inhibitors that block PMX activity do not inhibit its maturation (16). Therefore, we tagged PMX in the PfERC conditional mutants to test if it plays a role in PMX maturation. Our data show that PfERC knockdown inhibits the proteolytic maturation of PMX, suggesting that PfERC functions upstream to regulate the protease, directly or indirectly, responsible for PMX cleavage (Figure 7F). However, the protease responsible for PMX cleavage is unknown and there are no obvious candidates among the secreted proteases in the *Plasmodium* genome. Thus, in the absence of PfERC, the egress proteolytic cascade is inactive, which inhibits the proteolytic cleavage of PMX and SUB1. Our data suggest that PfERC does not play a role in signal-dependent vesicle exocytosis. Therefore, it is likely that uncleaved PMX and SUB1 are released from exonemes into the PV. However, since the unprocessed forms of PMX and SUB1 are inactive, they fail to breakdown the PVM as well as to cleave essential invasion ligands on the merozoite surface, such as MSP1 and AMA1.

A principal finding of these studies is the identification of an early regulator in the ER of *P. falciparum* with a specific role in egress of malaria parasites from RBCs and potentially in the invasion of parasites into the RBC. These data help build a model where PfERC modulates the maturation of the egress proteolytic cascade. The discovery of PfERC now provides us with a thread to unravel the rest of the egress proteolytic cascade in malaria parasites. These studies lay the foundation for understanding the vital and key role that ER-resident proteins play in the egress of human malaria parasites from the infected RBC and their re-entry into the host cell. Some studies have suggested that a key class of antimalarials containing endoperoxides, which includes the frontline antimalarial artemisinin, may target PfERC (46) and one of the transcriptomic responses of artemisinin-resistant parasites is the overexpression of PfERC (55). These data suggest that targeting PfERC, and thus egress, is a viable strategy for antimalarial drug development.

## Material and Methods

### Cell culture and transfections

*Plasmodium* parasites were cultured in RPMI 1640 medium supplemented with Albumax I (Gibco) and transfected as described earlier (56–59). To generate PfERC*-glmS* and PfERC*-M9* parasites, a mix of two plasmids (50µg of each) was transfected in duplicate into 3D7 parasites. The plasmid mix contained pUF1-Cas9-guide (60) which contains the DHOD resistance gene, and pPfERC-HA-SDEL-*glmS* or pPfERC-HA-SDEL-*M9*, which are marker-free. Drug pressure was applied 48hrs after transfection, using 1µM DSM1 (61), selecting for Cas9 expression. DSM1 was removed from the culturing medium once the parasites were detected in the culture, around 3 weeks post-transfection.

To generate PfERC-*glmS*/PMX-tetR and PfERC-*M9*/PMX-tetR parasites, we transfected PfERC-*glmS* and PfERC-*M9* parasites with a mix of two plasmids; pPMX-V5-Apt-pMG74 and the pyAIO plasmid containing the PMX guide RNA (16). Before transfection, 20µg of pPMX-V5-Apt-pMG74 was digested overnight with EcoRV. The enzyme was then heat inactivated for 20min at 65°C and DNA was co-precipitated with 50µg of PMX-guideRNA pyAIO. Transfected parasites were grown in 0.5 µM anhydrous tetracyline (aTc) (Cayman Chemical). Drug pressure was applied 48hrs after transfection, using BSD at a concentration of 2.5 µg/mL (61), selecting for pPMX-V5-Apt-pMG74 expression. Two clones each were isolated from the PfERC-*glmS* and the PfERC-*M9* backgrounds.

### Construction of PfERC and PMX plasmids

Genomic DNA was isolated from *P. falciparum* cultures using the QIAamp DNA blood kit (Qiagen). Constructs utilized in this study were confirmed by sequencing. PCR products were inserted into the respective plasmids using the In-Fusion cloning system (Clontech), the sequence- and ligation-independent cloning (SLIC) method (58, 59), T4-ligation (New England BioLabs), or site-directed mutagenesis using QuickChange (Agilent). To generate the pHA-SDEL-*glmS*/*M9* plasmid, primers 1+2 were used to add an SDEL sequence at the end of the HA tag in pHA-*glmS* and pHA-*M9* plasmids (58, 59).

For generating the PfERC-*glmS*/*M9* conditional mutants, pHA-SDEL-*glmS*/*M9* plasmid, consisting of two homology regions flanking the HA-SDEL tag and the *glmS* or *M9* sequence, was used as a donor DNA template. To allow efficient genomic integration of the pHA-SDEL-*glmS* and pHA-SDEL-*M9* donor plasmids, 800-bp sequences were used for each homology region. The C-terminus of the *pferc* coding region was PCR amplified from genomic DNA using primers 3+4 (containing the shield mutation) and was inserted into pHA-SDEL-*glmS* and pHA-SDEL-*M9* using restriction sites SacII and AfeI. The 3’UTR of *pferc* was PCR amplified from genomic DNA using primers 5+6 and was inserted into pHA-SDEL-*glmS* and pHA-SDEL-*M9* (already containing the C-terminus region) using restriction sites HindIII and NheI. For expression of PfERC guide RNA, oligos 7+8 were inserted into pUF1-Cas9-guide as previously described (58, 59). Briefly, pUF1-Cas9-guide was digested with BtgZI and annealed oligos were inserted using SLIC. Primers 3+6 and primers 3+9 (which recognizes the *glmS*/*M9* sequence) were used for clone verification.

To generate the pPMX-V5-Apt-pMG74, primers 10+11 were used to amplify the 3’UTR homology region containing AflII and EcoRV sites respectively and primers 12+13 were used to amplify the C-terminus homology region containing EcoRV and PspXI sites respectively. Then, using primers 10+13 the two PCR products stitched together. This final PCR product was ligated into pPMX-V5-Apt-pMG74 using AflII and PsPXI sites by SLIC.

### *Plasmodium* growth assays

Asynchronous growth assays were done as described previously (71, 72). Briefly, 5mM glucosamine (GlcN) (Sigma) was added to the growth medium and parasitemia was monitored every 24hrs using a CyAn ADP (Beckman Coulter) or CytoFLEX (Beckman Coulter) flow cytometers and analyzed by FlowJo software (Treestar, Inc.). As required, parasites were subcultured to avoid high parasite density, and relative parasitemia at each time point was back-calculated based on actual parasitemia multiplied by the relevant dilution factors. One hundred percent parasitemia was determined as the highest relative parasitemia and was used to normalize parasite growth. Data were fit to exponential growth equations using Prism (GraphPad Software, Inc.).

To determine the ring:schizont ratio of PfERC-*glmS* and PfERC-*M9* parasites, 7.5mM GlcN was added to percoll isolated schizont-stage parasites and parasites were allowed to egress and reinvade fresh RBCs. Two hours later, 5% sorbitol +7.5mM GlcN was added to the invaded culture to lyse any remaining schizonts and isolate two-hour rings. The ring-stage parasites were grown again in media supplemented with GlcN. Then samples were taken at 44hrs, 48hrs, and 56hrs, and read by flow cytometry to determine the population of rings and schizonts present at those times using FlowJo software (Treestar, Inc.). To determine the development of each life cycle stage during the asexual lifecycle of PfERC-*glmS* and PfERC-*M9* parasites, 7.5mM was added to percoll isolated schizont-stage parasites and parasites were allowed to egress and reinvade fresh RBCs. At specific times Hema-3 stained blood smears were used to count parasite stages and the percentage of the specific lifecycle stage was calculated as: 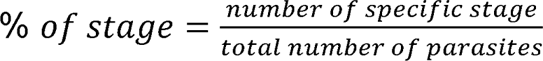. Time 0hr is when GlcN was added.

To determine the % amount of rings or schizonts, samples of synchronized schizonts grown with 7.5mM GlcN for about 48hrs were taken and fixed with 8% paraformaldehyde and 0.3% glutarladehyde. Samples were read by flow cytometry. For growth assays using Compound 1 (4-[2-(4-fluorophenyl)-5-(1-methylpiperidine-4-yl)-1H-pyrrol-3-yl] pyridine), synchronized schizonts were grown with 7.5mM GlcN for about 48hrs. Then, schizonts were percoll isolated and incubated with Compound 1 for 4hrs and then removed by gently washing parasites twice with 1mL of warm, complete RPMI + 7.5mM GlcN. Parasites were resuspended with fresh media and RBCs and fixed samples (as above) were read by flow cytometry. DNA content was determined using Hoechst 33342 staining (ThermoFisher).

### Western blotting

Western blotting for *Plasmodium* parasites was performed as described previously (58, 59). Briefly, parasites were permeabilized selectively by treatment with ice-cold 0.04% saponin in PBS for 10 min and pellets were collected for detection of proteins with the parasite. For detection of MSP1, schizonts were isolated on a Percoll gradient (Genesee Scientific) and whole-cell lysates were generated by sonication. The antibodies used in this study were rat anti-HA (3F10; Roche, 1:3,000), rabbit anti-HA (715500; Invitrogen, 1:100), mouse anti-V5 (TCM5; eBioscience, 1:1000), rabbit anti-PfEF1α (from D. Goldberg, 1:2,000), mouse anti-plasmepsin V (from D. Goldberg, 1:400), rabbit anti-SUB1 (from Z. Dou and M. Blackman, 1:10,000), rat anti-AMA1 (28G2; Alan Thomas via BEI Resources, NIAID, NIH 1:500), mouse anti-MSP1 (12.4; European Malaria Reagent Repository, 1:500) and mouse anti-RAP1 (2.29; European Malaria Reagent Repository, 1:500). The secondary antibodies that were used are IRDye 680CW goat anti-rabbit IgG and IRDye 800CW goat anti-mouse IgG (LICOR Biosciences) (1:20,000). The Western blot images were processed using the Odyssey Clx LICOR infrared imaging system software (LICOR Biosciences). Calculation of knockdown and processing ratios was determined by both the Odyssey infrared imaging system software and ImageJ 1.8 (NIH).

### Immunofluorescence microscopy

For IFAs, cells were fixed as described previously (58, 59). The antibodies used for IFA were: rat anti-HA antibody (clone 3F10; Roche, 1:100), mouse anti-AMA1 (1F9 from Alan Cowman), rat anti-PfGRP78 (MRA-1247; BEI Resources, NIAID, NIH 1:100), mouse anti-MSP1 (12.4; European Malaria Reagent Repository, 1:500), rat anti-AMA1 (28G2; Alan Thomas via BEI Resources, NIAID, NIH 1:500), and mouse anti-RAP1 (2.29; European Malaria Reagent Repository, 1:500). Secondary antibodies used were anti-rat antibody conjugated to Alexa Fluor 488 or 546 and anti-rabbit antibody conjugated to Alexa Fluor 488, (Life Technologies, 1:100). Cells were mounted on ProLong diamond with 4’,6’-diamidino-2-phenylindole (DAPI) (Invitrogen) and imaged using a Delta-Vision II microscope system with an Olympus IX-71 inverted microscope using a 100x objective or an Elyra S1 SR-SIM microscope (Zeiss). Image processing, analysis, and display were performed using SoftWorx or Axiovision and Adobe Photoshop. Adjustments to brightness and contrast were made for display purposes.

### AMA1 Translocation Assays

To observe AMA1 translocation in our mutants, 7.5mM GlcN was added to percoll isolated schizont-stage parasites and parasites were allowed to egress and reinvade fresh RBCs. 44-48hrs later, schizonts were percoll purified and incubated with 1.5µM Compound 1 for 4 hours at 37°C. Then, Compound 1 removed by washing parasites twice with 1mL of warm complete RMPI +7.5mM GlcN. These parasites were immediately resuspended in media plus 7.5mM GlcN and 20 µM E-64 (Sigma) and incubated at 37°C in a still incubator for 6hrs. Parasites were then fixed as in (58, 59) and probed with anti-AMA1 (1F9) antibodies. Images were taken using a Delta-Vision II microscope system with an Olympus IX-71 inverted microscope using a 100x objective and using an Elyra S1 SR-SIM microscope (Zeiss).

### Immunoprecipitation Assay

For IP, we took samples of percoll purified schizonts form PfERC-*M9* parasites. Collected schizonts were incubated on ice for 15min in Extraction Buffer (40mM Tris-HCl, 150mM KCl, 1M EDTA plus 1XHALT and 0.5% NP40). The parasites were lysed via sonication and the sample was centrifuged at 20,000 g in 4°C for 15 min. The supernatant was mixed with anti-HA antibody-conjugated magnetic beads (Thermo Scientific) for 1hr at room temperature. The beads were washed 3X with IgG Binding Buffer (20mM Tris-HCl, 150mM KCl, 1mM EDTA, 0.1% NP40) and, elution was done with 100uL of 2mg/mL HA peptide (Thermo Scientific) at 37°C for 10 minutes while rocking.

### Transmission Electron Microscopy

7.5mM GlcN was added to percoll isolated schizont-stage parasites and parasites were allowed to egress and reinvade fresh RBCs. 48hrs later, parasites were percoll-isolated and then incubated with 20µM E-64 for 8hrs. After incubation, parasites were washed with 1X PBS and gently resuspended in 2.5% glutaraldehyde in 0.1M sodium cacodylate-HCl (Sigma) buffer pH 7.2 for 1hr at room temperature. Parasites were then rinsed in 0.1M Cacodylate-HCl buffer before agar-enrobing the cells in 3% Noble agar. Parasites were post fixed in 1% osmium tetroxide/0.1M Cacodylate-HCl buffer for 1 hour and rinsed in buffer and deionized water. Dehydration of the parasite samples was done with an ethanol series and then exposed to Propylene oxide before infiltration with Epon-Araldite. The blocks of parasites were trimmed, and sections were obtained using a Reichert Ultracut S ultramicrotome (Leica, Inc., Deerfield, IL). 60-70nm sections were placed on 200-mesh copper grids and post-stained with ethanolic uranyl acetate and Reynolds Lead Citrate. Grids were viewed with a JEOL JEM-1011 Transmission Electron Microscope (JEOL USA, Inc., Peabody, MA) using an accelerating voltage of 80 KeV. Images were acquired using an AMT XR80M Wide-Angle Multi-Discipline Mid-Mount CCD Camera (Advanced Microscopy Techniques, Woburn, MA).

### Scanning Electron Microscopy

7.5mM GlcN was added to percoll isolated schizont-stage parasites and parasites were allowed to egress and reinvade fresh RBCs. 48hrs later, parasites were percoll-isolated and then incubated with 2µM Compound 2 (4-[7-[(dimethylamino)methyl]-2-(4-fluorphenyl)imidazo[1,2-a]pyridine-3-yl]pyrimidin-2-amine) for 4 hours without shaking at 37°C in an incubator. After incubation, parasites were washed twice with warm, complete RPMI + 7.5mM GlcN. Samples were taken immediately after washing off Compound 2 and then 30min after and fixed as with TEM samples. Parasites were rinsed with 0.1M Cacodylate-HCl buffer before placing on glass coverslips prepared with 0.1% Poly-L-lysine. Parasites were allowed to settle onto the glass coverslips in a moist chamber overnight and then post fixed in 1% osmium tetroxide/0.1M Cacodylate-HCl buffer for 30 minutes. Cells on coverslips were rinsed three times in deionized water and then dehydrated with an ethanol series. The glass coverslips were critical point dried in an Autosamdri-814 Critical Point Dryer (Tousimis Research Corporation, Rockville, MD), mounted onto aluminum pin stubs with colloidal paint, and sputter coated with gold-palladium with a Leica EM ACE600 Coater (Leica Microsystems Inc., Buffalo Grove, IL). Stubs were examined with the FE-SEM FEI Teneo (FEI, Inc., Hillsboro, OR) using the secondary electron detector to obtain digital images.

### Calcium Measurements

To measure Ca^2+^ in PfERC mutants, knockdown was induced on synchronized schizonts. After 48hrs, schizonts were percoll purified and permeabilized selectively by treatment with ice-cold 0.04% saponin in PBS for 10 min. Isolated parasites were then washed 2X with BAG Buffer (116mM NaCl, 5.4mM KCl, 0.8mM MgSO_4_·7H_2_O, 50mM HEPES, 5.5mM Glucose) + 7.5mM GlcN and incubated with 10µM Fluo-4AM (ThermoFisher) for 45min while rocking at 37°C. After incubation, cells were washed 2X with BAG buffer + 7.5mM GlcN and immediately taken for fluorimetric measurements. Fluorescence measurements were carried out in a cuvette (Sarstedt) containing parasites suspended in 2.5 ml of BAG buffer and 100uM EGTA (Sigma). The cuvette was placed in a Hitachi F-4500 fluorescence spectrophotometer and Fluo-4AM excitation was done at 505 nm with emission read at 530 nm (63). Drugs and reagents were added via a Hamilton syringe. Final concentration of CPA (Sigma) was 3 µM, and Ionomycin (Sigma) at 2µM.

## Acknowledgments

We thank Michael Reese and Meg Phillips for comments on the manuscript; Dan Goldberg for comments on the manuscript, anti Ef1α and anti-PMV antibodies, and the pyAIO plasmid; The European Malaria Reagent Repository for anti-MSP1 12.4 and anti-RAP1 2.29, antibodies; Alan Thomas and BEI Resources NIAID, NIH for anti-AMA1 28G2 and anti-BiP antibodies; Zhicheng Dou and Michael Blackman for anti-SUB1 antibody; Alan Cowman for anti-AMA1 antibody (1F9); Purnima Bhanot for Compound 1 and Compound 2; Muthugapatti Kandasamy at the University of Georgia Biomedical Microscopy Core, Julie Nelson at the CTEGD Cytomtetry Shared Resource Lab for technical assistance; Mary Ard from the Georgia Electron Microscopy core at the University of Gerogia for assistance with SEM and TEM; Michael Cipriano for assistance with protein alignment. This work was supported by UGA Faculty Research Grant (FRG-SE0031) to V.M., and the US National Institutes of Health (R21AI133322) to V.M. and S.N.J.M., and (T32AI060546) to M.A.F. and D.W.C.

---

**Table S1:**
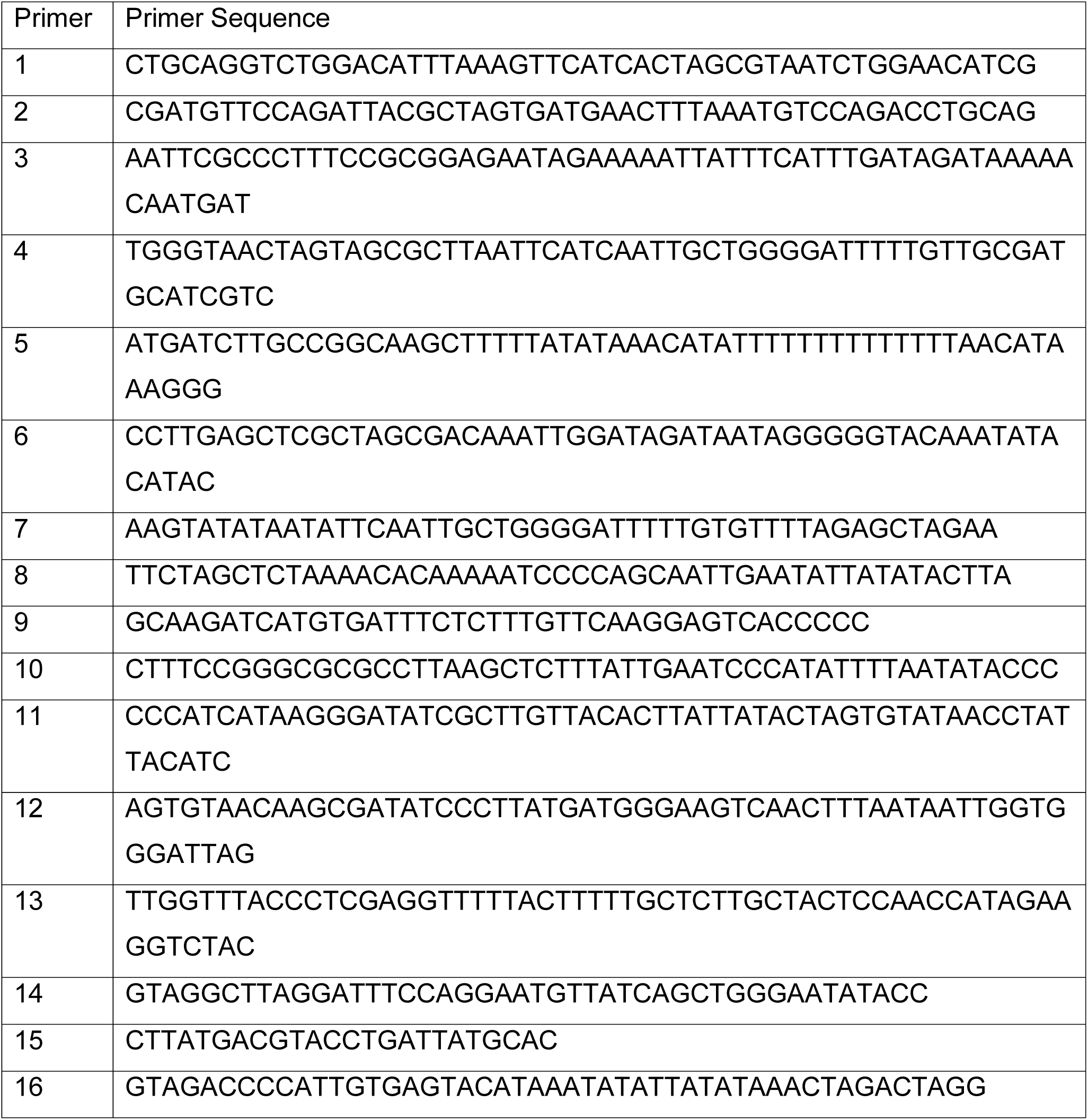
Primers used in this study.

**Supplementary Figure 1:**
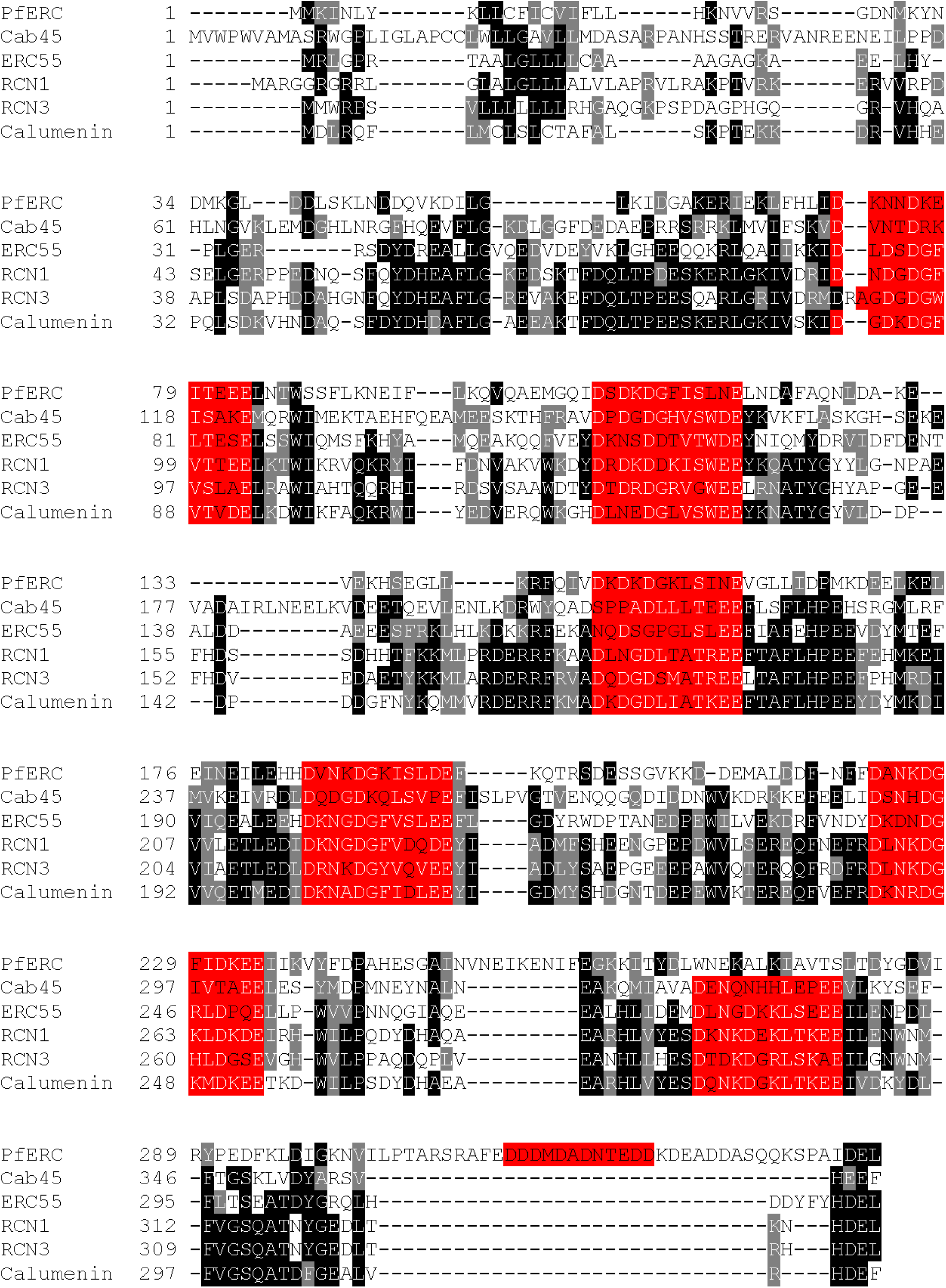
Sequence alignment of PfERC to other members of the CREC family of proteins using MUSCLE alignment, viewed using JalView Software (http://www.jalview.org/) and BOXSHADE (82). Alignment was done using the human homologs: Cab-45, ERC-55, Reticulocalbin1(RCN1), Reticulocalbin (RCN3), and Calumenin. Identical residues are shaded in black, similar residues are shaded in gray, and EF-hands are highlighted in red.

**Supplementary Figure 2:**
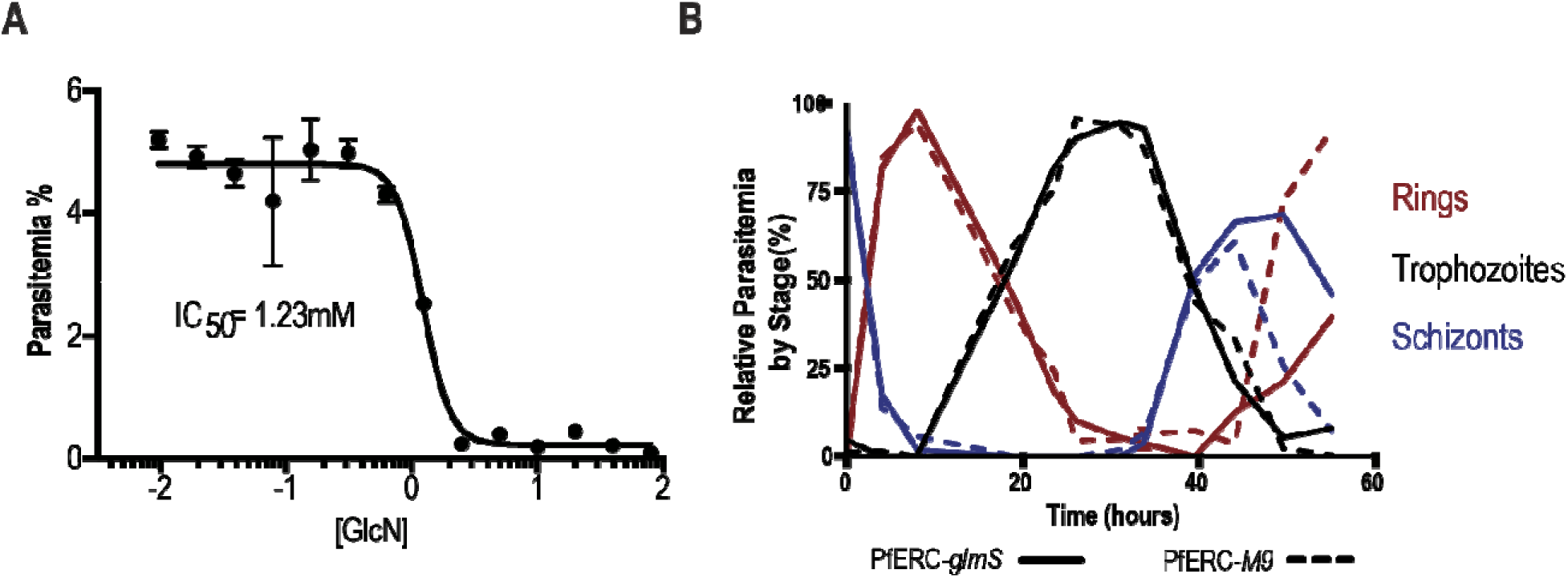
Effect of PfERC knockdown on parasite growth. (A) Asynchronous PfERC-*glmS* parasites were incubated in different concentrations of GlcN and growth after three days was assessed by flow cytometry. Data are represented as mean ± SEM of n=2 biological replicates. (B) The levels of rings (red), trophozoites (black), and schizonts (blue) as a percentage of total parasites as scored by light microscopy of Hema-3 stained blood smears from synchronous PfERC-*glmS* and PfERC-*M9* parasites grown in the presence of GlcN (n=2 biological replicates).

**Supplementary Figure 3:**
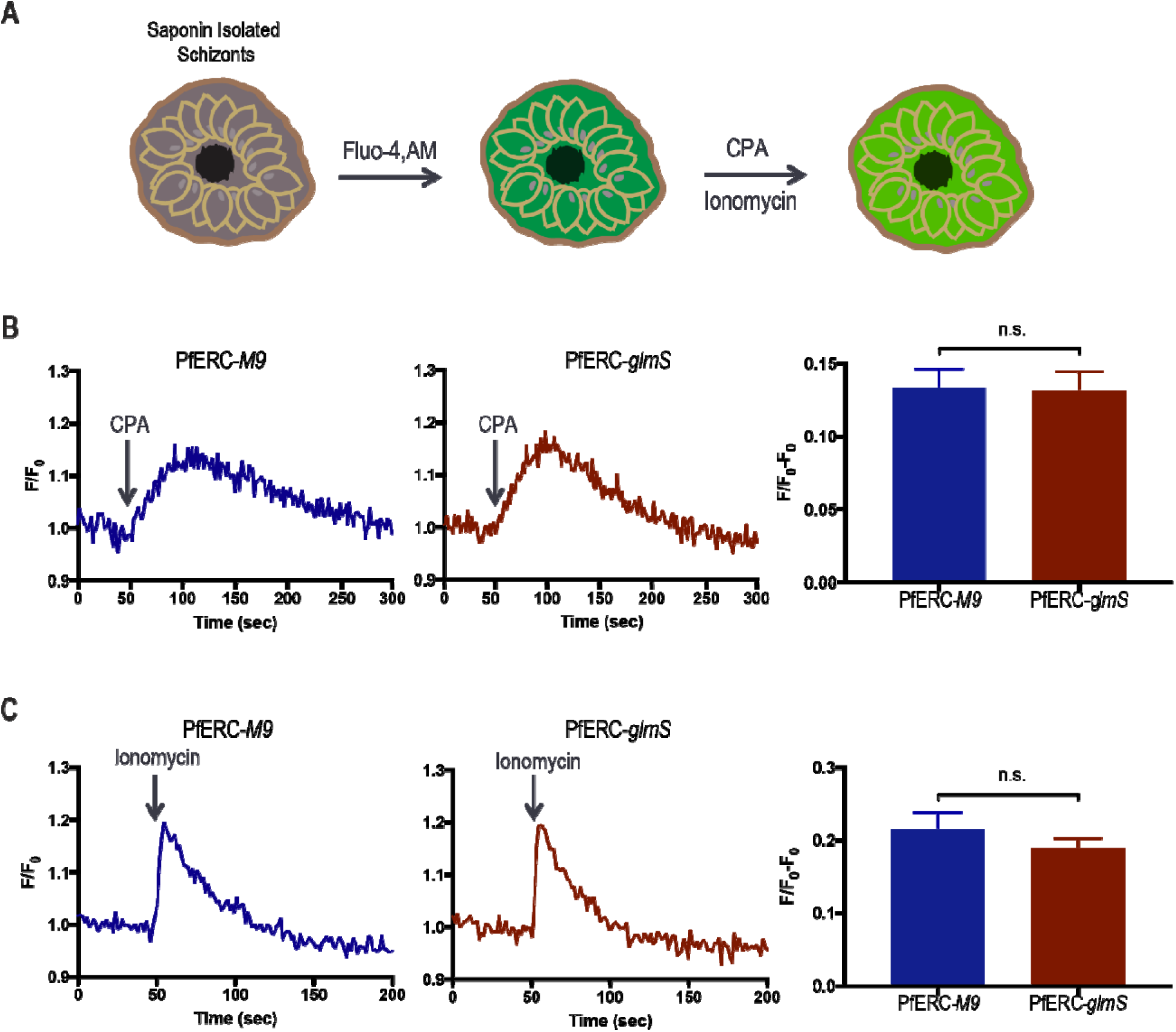
PfERC knockdown is not required for ER Ca^2+^ storage. (A) Experimental schematic showing how Ca^2+^ measurements were done in PfERC-*glmS* and PfERC-*M9* mutants. Synchronized PfERC-*glmS* and PfERC-*M9* schizonts were incubated with GlcN for 48 hours and isolated using saponin lysis, which lyses the RBC membrane but leaves the PV intact. Abbreviations: CPA-cyclopiazonic acid (B) Representative fluorescence tracings after CPA addition to PfERC-*glmS* and PfERC-*M9* schizonts, isolated as in (A). Quantification was done by calculating the difference in fluorescence between the basal to the highest peak of fluorescence. Data are represented as the combined mean ± SEM (PfERC-*glmS;* n=15 biological replicates; PfERC-*M9;* n=9 biological replicates; n.s-non-significant, unpaired *t-*test). (C) Representative fluorescence tracings after Ionomycin addition to PfERC-*glmS* and PfERC-*M9* schizonts, isolated as in (A). Quantification was done by calculating the difference in fluorescence between the basal to the highest peak of fluorescence. Data are represented as the combined mean ± SEM (PfERC-*glmS;* n=9 biological replicates; PfERC-*M9;* n=5 biological replicates; n.s-non-significant, unpaired *t-*test).

**Supplementary Figure 4:**
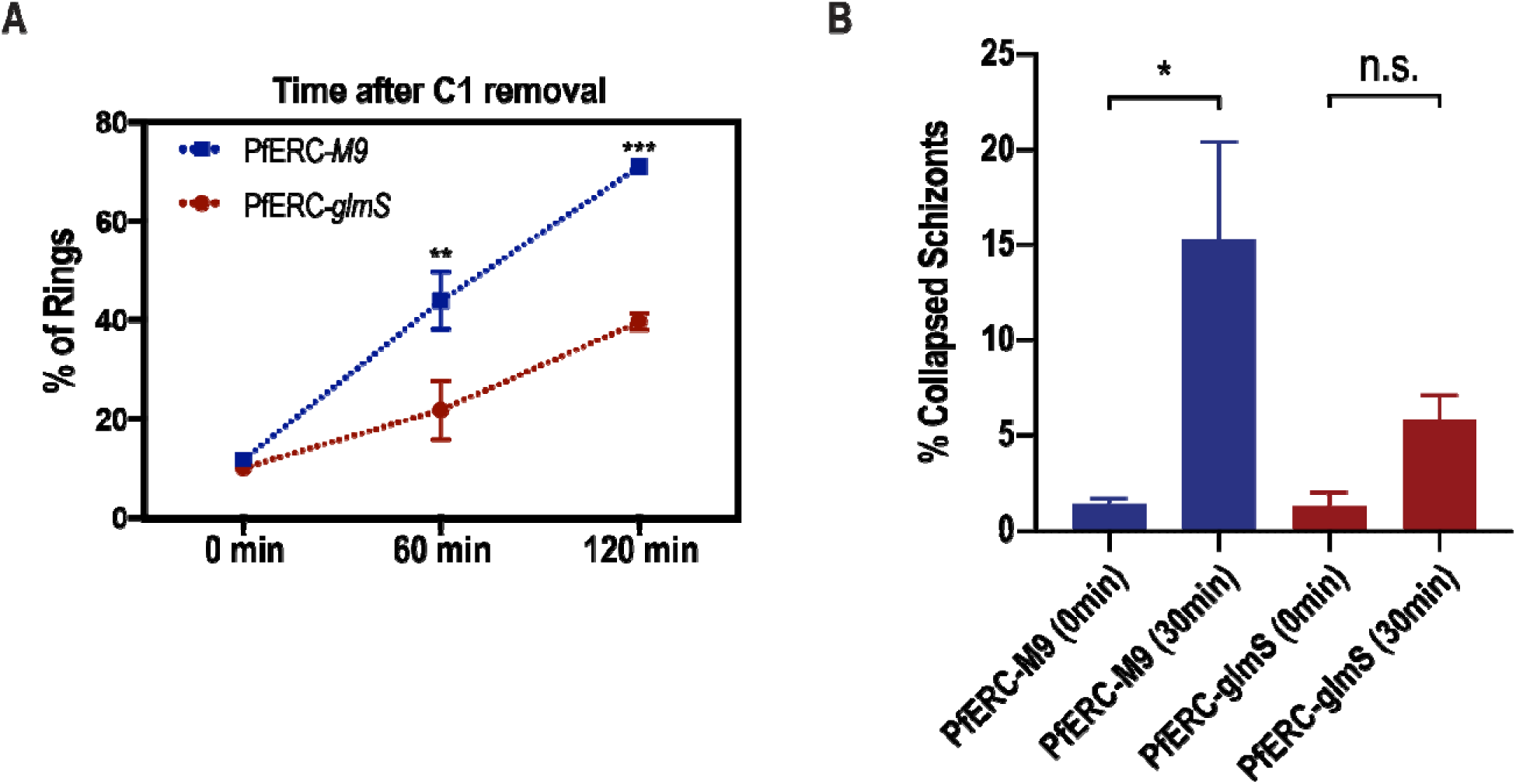
A) As shown in Figure 3A, synchronized PfERC-*glmS* and PfERC-*M9* schizonts were grown in the presence of GlcN and second cycle rings were observed by flow cytometry after removal of C1 (time 0hr). Rings were quantified as a percentage of the total amount of parasites as determined by flow cytometry. Data are represented as the mean ± SEM (n=3 biological replicates; **P<0.01, ***P<0.001 2-way ANOVA). B) PfERC-*glmS* and PfERC-*M9* schizonts were treated as shown in Figure 3C and wide field (10 fields per biological replicate) SEM images were quantified. The collapsed schizonts as shown in Figure 3C were normalized to total schizonts counted in the fields. Data are represented as mean ± SEM (n=4 biological replicates; n.s.=non-significant, \**P*<0.05 one-way ANOVA).

**Supplementary Figure 5:**
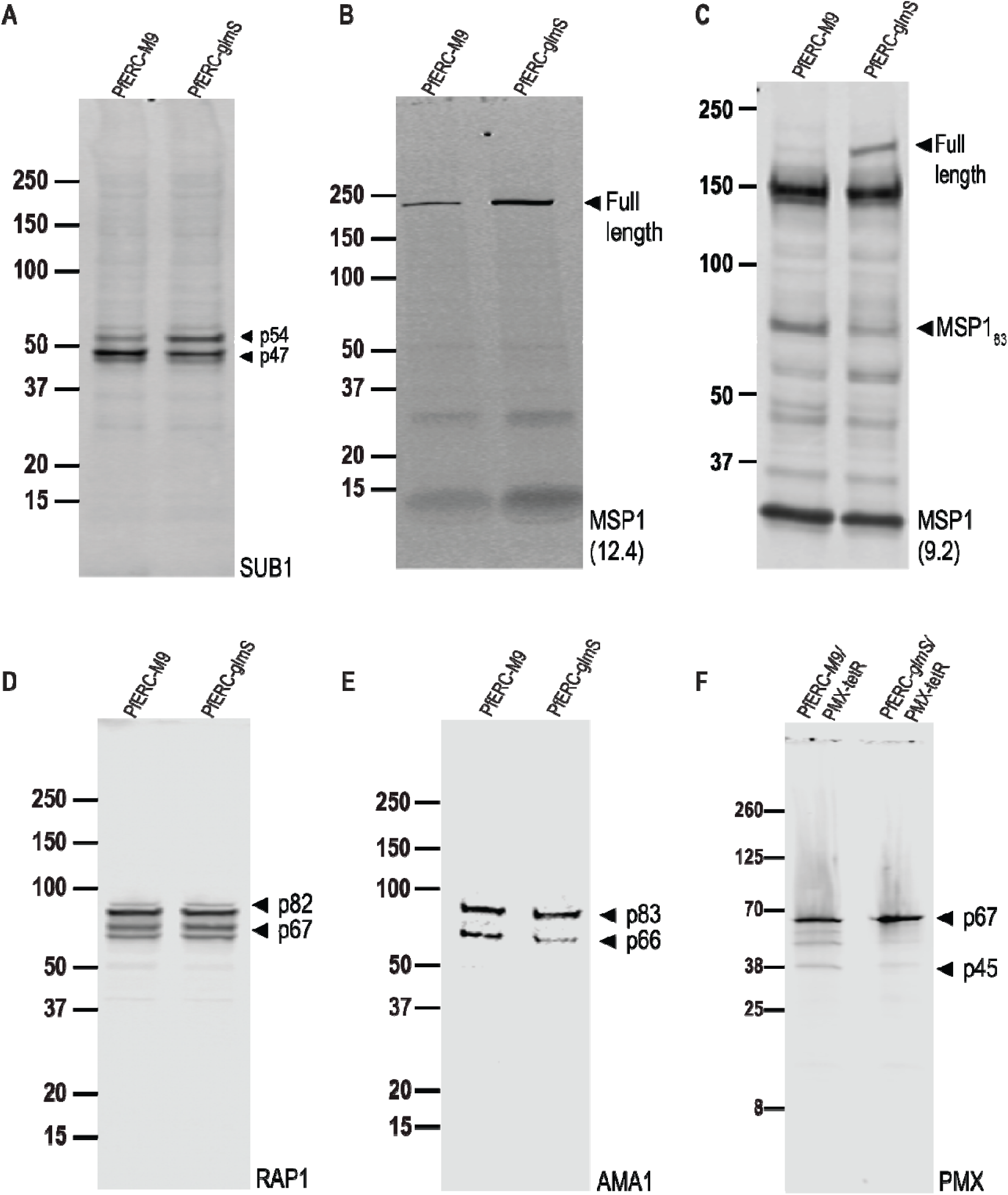
Representative images of Western blots of lysates from PfERC-*glmS* and PfERC-*M9* schizonts incubated with GlcN for 48 hours, probed with anti-SUB1, anti-MSP1 12.4 and 9.2, anti-AMA1, anti-RAP1, and anti-V5 antibodies from Figures 4, 5, 6 and 7. The protein marker sizes that co-migrated with the probed protein are shown on the left.

**Supplementary Figure 6:**
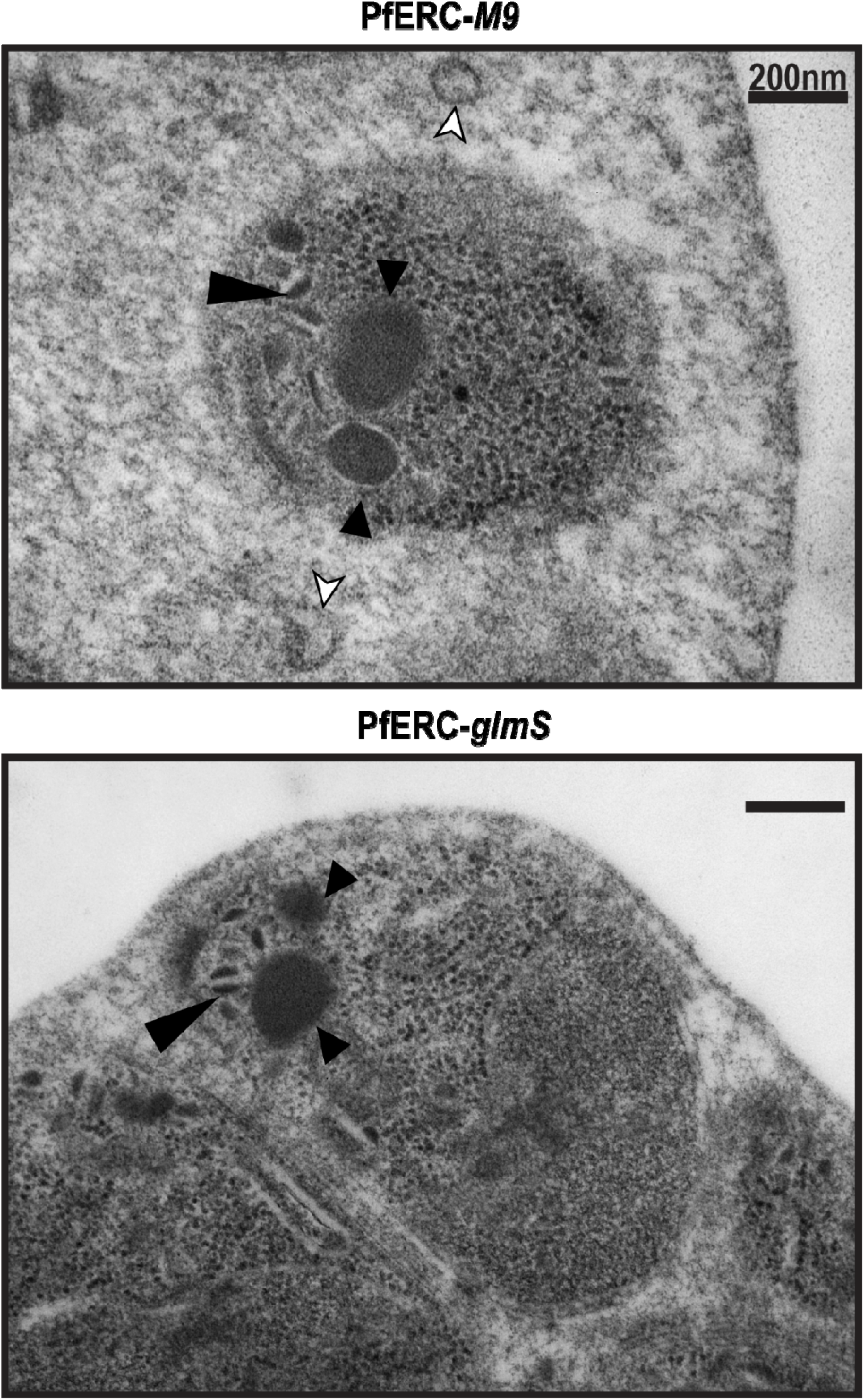
Representative TEM images of PfERC-*glmS* and PfERC-*M9* schizonts grown for 48hrs with GlcN and incubated with E-64 for 8 hours, as shown in Figure 3A (n=2 biological replicates). Small arrowheads point to rhoptries, large arrowheads to micronemes, and white arrowheads to PVM fragments (15). Scale bar, 200nm.

**Supplementary Figure 7:**
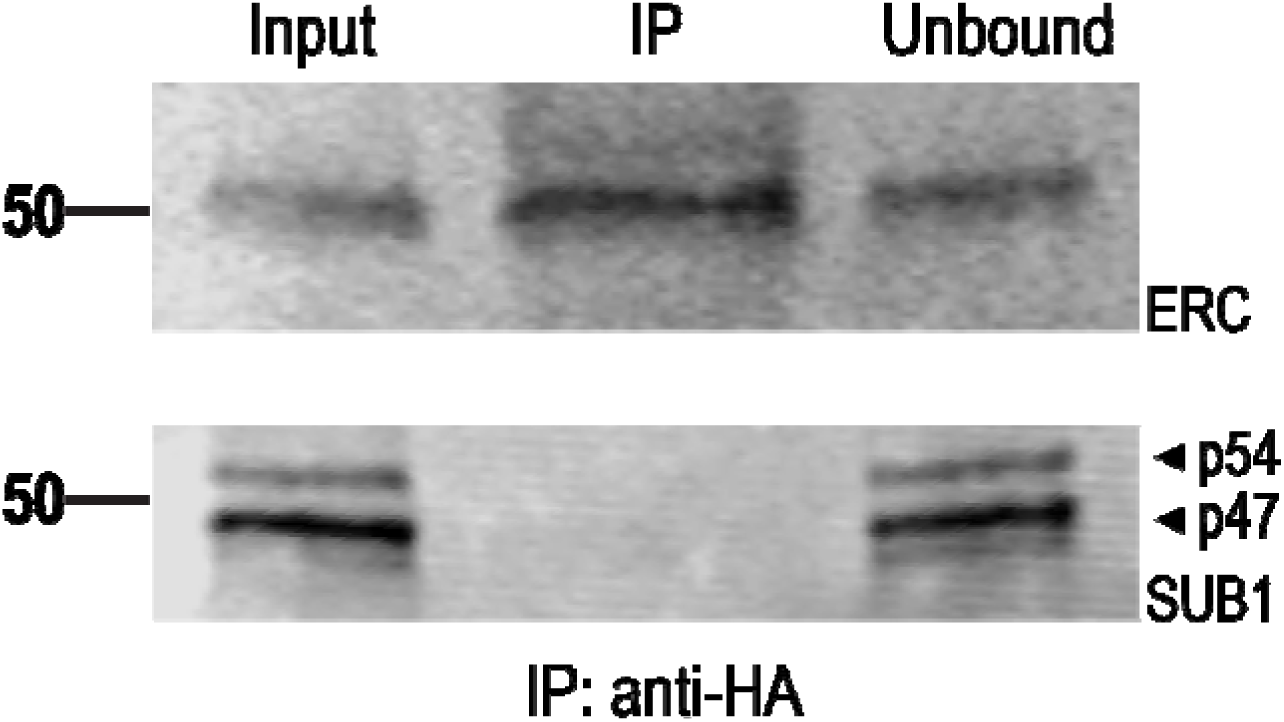
Co-Immunoprecipitation of PfERC and SUB1. PfERC-*M9* schizonts were isolated and immunoprecipitated using HA-conjugated beads. The input, IP, and unbound fractions were probed with anti-HA and anti-SUB1 antibodies (n=2 biological replicates).

